# Sustainable Production of the Biofuel *n*-Butanol by *Rhodopseudomonas palustris* TIE-1

**DOI:** 10.1101/2020.10.13.336636

**Authors:** W. Bai, T. O. Ranaivoarisoa, R. Singh, K. Rengasamy, A. Bose

## Abstract

Anthropogenic carbon dioxide (CO_2_) release in the atmosphere from fossil fuel combustion has inspired scientists to study CO_2_ to fuel conversion. Oxygenic phototrophs such as cyanobacteria have been used to produce biofuels using CO_2_. However, oxygen generation during oxygenic photosynthesis affects biofuel production efficiency. To produce *n*-butanol (biofuel) from CO_2_, here we introduced an *n*-butanol biosynthesis pathway into an anoxygenic (non-oxygen evolving) photoautotroph, *Rhodopseudomonas palustris* TIE-1 (TIE-1). Using different carbon, nitrogen, and electron sources, we achieved *n*-butanol production in wild-type TIE-1 and mutants lacking electron-consuming (nitrogen-fixing) or acetyl-CoA-consuming (polyhydroxybutyrate and glycogen synthesis) pathways. The mutant lacking the nitrogen-fixing pathway produced highest *n*-butanol. Coupled with novel hybrid bioelectrochemical platforms, this mutant produced *n*-butanol using CO_2_, solar panel-generated electricity, and light, with high electrical energy conversion efficiency. Overall, this approach showcases TIE-1 as an attractive microbial chassis for carbon-neutral *n*-butanol bioproduction using sustainable, renewable, and abundant resources.

## Introduction

The rapid consumption of fossil fuels has increased carbon dioxide (CO_2_) levels in the atmosphere raising concerns about global warming^1,2^. This environmental concern has spurred research initiatives aiming to develop carbon-neutral biofuels that, when burned, will not result in net CO_2_ release^3^. Among the various biofuels, *n*-butanol has received greater attention due to its higher energy content, lower volatility, and reduced hydrophilicity compared to ethanol^4^. Currently, most *n*-butanol is synthesized via chemical processes^5,6^. However, these processes use propylene or ethanol as feedstock, making these methods, carbon-positive^5,6^. Another well-known strategy for *n*-butanol production is the acetone–butanol–ethanol (ABE) fermentation using *Clostridium* species^7^. As such, the *n*-butanol biosynthesis pathway^7^ (Fig. 1a) from *Clostridium acetobutylicum* has been introduced into several organisms, such as *Escherichia coli, Saccharomy*ces *cerevisiae, Pseudomonas putida*, and *Bacillus subtilis* for *n*-butanol production^8–11^. However, most of these organisms are chemoheterotrophs. Thus, the *n*-butanol production using these microbes is also carbon-positive.

**Figure 1.**
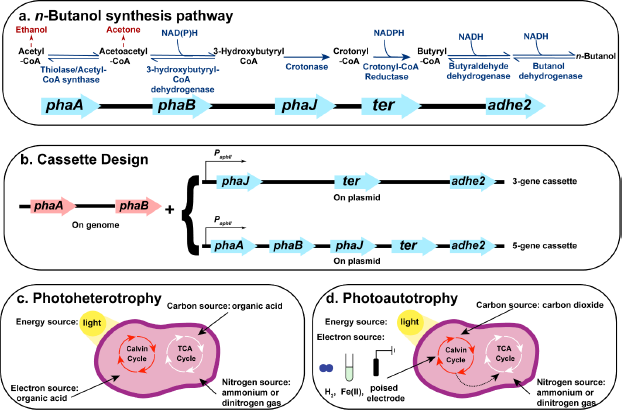
*n*-Butanol synthesis pathway, cassette design and major metabolisms used for *n*-butanol production in *Rhodopseudomonas palustris* TIE-1. (**a**) *n*-butanol biosynthesis pathway involves five genes. The enzymes encoded by each gene and the reactions catalyzed by these enzymes are shown in dark blue. Two major byproducts (acetone and ethanol) are shown in dark red. NADH, nicotinamide adenine dinucleotide. (**b**) Cassette design. The 3-gene cassette relies on *phaA* and *phaB* on the genome of TIE-1 for the first two steps of *n*-butanol synthesis. Here, only 3-genes (*phaJ, ter*, and *adhE*2) were introduced on a plasmid under a constitutive promoter. The 5-gene cassette has all five genes (*phaA, phaB, phaJ, ter*, and *adhE*2) on the plasmid under a constitutive promoter (**c**) Photoheterotrophy: TIE-1 uses organic acids as carbon and electron source, light as an energy source, and ammonium or dinitrogen gas as a nitrogen source. (**d**) Photoautotrophy: TIE-1 uses carbon dioxide (CO_2_) as carbon source, hydrogen (H_2_), ferrous iron (Fe(II)), or poised electrode as an electron source, light as an energy source, and ammonium or dinitrogen gas as a nitrogen source

To date, only a handful of studies have produced *n*-butanol autotrophically using CO_2_ as a carbon source^12–16^. Using a microbial electrosynthesis approach, a chemoautotroph *Clostridium* sp. produced 135 mg/L of *n*-butanol at an applied potential (E_appl_) of 0.8 V using CO_2_ in 35 days^12^. This prolonged period was required for acid accumulation for *n*-butanol production using *Clostridium* sp.^7^. Autotrophic *n*-butanol production was also demonstrated by an oxygenic photoautotroph *Synechococcus elongatus* PCC 7942 using water as an electron donor^14^ and sunlight as the energy source. Because, the *n*-butanol was generated using solar energy, this product is called a solar fuel. With the *n*-butanol biosynthesis pathway, *S. elongatus* produced 2.2 mg/L *n*-butanol^14^ when incubated anaerobically under illumination. In contrast, aerobic incubation did not generate any *n*-butanol. Furthermore, a dark anaerobic incubation of dense cultures (where cells were not actively growing or evolving oxygen) produced 14 mg/L of *n*-butanol^14^. These results suggest that oxygen (O_2_) is detrimental to *n*-butanol production^14^. The ability of cyanobacteria to produce *n*-butanol was later improved by several modifications such as 1) using cofactor as a driving force^15^; 2) replacing the oxygen-sensitive enzyme involved in the *n*-butanol producing pathway^16^, and 3) using intensive engineering to optimize the pathway in a multi-level modular manner^17^. These engineered cyanobacterial strains produced 29.9 mg/L^15^, 404 mg/L^16^, and 4.8 g/L^17^ of *n*-butanol. However, the low (<3%) energy conversion efficiency of natural photosynthesis^18^ makes the use of cyanobacteria not ideal for *n*-butanol production.

To enhance energy conversion efficiency for biofuel production, artificial photosynthesis, where photo-generated electrons were used to drive chemical reactions^19^, was developed. However, due to catalyst limitations, hydrogen (H_2_) was the main product^20^. Although H_2_ can be used as a fuel, using such an explosive gas requires significant modifications to the current gasoline-based infrastructures^21,22^. To avoid this, a H_2_-consuming chemoautotrophic bacterium *Ralstonia eutropha*, was used for producing carbon-based liquid fuels using hybrid water-splitting biosynthetic system. In this system, H_2_ and O_2_ were produced from water splitting (powered by electricity from a potentiostat) using a cobalt phosphorus catalyst with an applied electrical potential (E_appl_) of 2.0 V^19^. The H_2_ was then fed to the engineered *R. eutropha* to synthesize C_3_-C_5_ alcohol or polyhydroxybutyrate (PHB) from CO_2_ ^19^. This hybrid biosynthetic system reached an electrical energy conversion efficiency (EECE) up to ~20% using air (400 ppm CO_2_) toward biomass^19^. These values far exceeded the energy conversion efficiency of natural photosynthesis Also, the system reported an EECE of 16 ± 2% towards C_4_~C_5_ alcohol using pure CO_2_. Coupling a solar panel/photovoltaic cell resulted in an energy conversion efficiency of 6% towards biomass with pure CO_2_^23^. Although these studies provided a platform for indirect solar production from CO_2_, this technology is not an efficient and economical method for biofuel synthesis because, 1) it produces O_2,_ which is detrimental to many biofuel synthesis processes^14^; 2) it uses H_2_ as an electron donor, which due to its low solubility limits electron transfer efficiency^24^ and, finally; 3) this system requires electrical potentials higher than 1.23 V^18^, making it an expensive method. Therefore, it is critical to look for organisms that can overcome these limitations to advance carbon-neutral biofuel production.

One such organism is the anoxygenic photoautotroph *Rhodopseudomonas palustris* TIE-1 (TIE-1). TIE-1 can use various carbon sources, such as atmospheric CO_2_ and organic acids that can be easily obtained from organic wastes^25^. TIE-1 can also fix dinitrogen gas (N_2_)^26^, and use a wide range of electron sources. These include H_2,_ which is a byproduct of many industries; ferrous iron [Fe(II)], which is a naturally abundant element^27,28^; and most importantly, poised electrodes (i.e., photoelectroautotrophy) for its photosynthetic growth^27,29–32^. This wide electron donor selection enables TIE-1 to perform photosynthesis while avoiding O_2_ generation, a harmful component for biofuel synthesis^14^. TIE-1’s ability to perform photoelectroautotrophy is advantageous for biofuel production because of the direct electron uptake by TIE-1 from a poised electrode bypasses the need for an indirect electron donor such as H_2_. TIE-1 has a low E_appl_ (0.1 V)^27,29–32^ requirement, which lowers cost and electrochemical O_2_ generation. TIE-1’s E_appl_ requirement is ~90% lower than that needed for water-splitting^19^, allowing us to use low-cost solar panels to build novel biohybrid systems for solar fuel synthesis. Overall, TIE-1 is a superlative biocatalyst that allows us to use extant CO_2_, N_2_, solar energy, and electrons generated by renewable electricity for bioproduction. This process enables excess electricity to be stored as a usable fuel or products for later use.

In a previous study *R. palustris* CGA009 (CGA009), a strain closely related to TIE-1, was engineered to produce *n*-butanol from *n*-butyrate^33^. In this study, the gene encoding the alcohol/aldehyde dehydrogenases (AdhE2) from *R. palustris* Bisb18 was codon-optimized and introduced into *R. palustris* CGA009^33^. When cultured in the absence of CO_2_, the modified CGA009 was forced to reduce *n*-butyrate into *n*-butanol to maintain the redox balance^33^. Although this study used a phototroph that is closely related to TIE-1 for *n*-butanol production, this approach is carbon-positive as it uses an organic substrate (*n*-butyrate).

To produce *n*-butanol in a sustainable and carbon-neutral manner, we introduced an efficient, codon-optimized *n*-butanol biosynthesis pathway into TIE-1. This pathway, was assembled using irreversible and efficient enzymes and produced 4.6 g/L *n*-butanol in *E. coli*^34^. The pathway contains five genes (*phaA, phaB, phaJ, ter, adhE*2)^27^. Because TIE-1 possesses homologs for the first two genes (*phaA* and *phaB*)^27^, we designed two different cassettes (Fig. 1b), containing either the whole (5-gene cassette) or a partial *n*-butanol biosynthesis pathway (3-gene cassette). As shown in Fig. 1a, carbon (acetyl-CoA), and reducing equivalents (NADH) are two major substrates for *n*-butanol biosynthesis. Previous studies in cyanobacteria have shown that a PHB synthase deletion mutant produces higher butanol^35^, and a glycogen synthase deletion mutant showed higher carbon conversion efficiency (CCE) towards iso-butanol^36^. We, therefore, constructed TIE-1 knockout mutants lacking hydroxybutyrate polymerase^18^ or glycogen synthase^37^. Previous studies suggested that nitrogenase deletion mutants possess a more reduced intracellular environment in *Rhodobacter capsulatus and* CGA009^37–39^. We predicted this would be true in TIE-1 and created a nitrogenase double mutant. After introducing the 3-gene cassette/5-gene cassette into the TIE-1 wild type (WT) and mutant strains, we tested *n*-butanol production under both photoheterotrophic (Fig. 1c) and photoautotrophic (Fig. 1d) conditions. Under photoelectroautotrophy, we used a novel hybrid bioelectrochemical cell (BEC) platform powered by electricity from either an electrical grid powered potentiostat or a solar panel.

Our results show that the anoxygenic phototroph TIE-1 can produce *n*-butanol sustainably using organic acids or CO_2_ as carbon source, light as an energy source, and H_2_, Fe(II), or electrons from renewably generated electricity as an electron source. To the best of our knowledge, this study represents the first attempt for biofuel production using a solar panel powered microbial electrosynthesis platform, where CO_2_ is directly converted to liquid fuel. Overall, these results show that TIE-1 can be an attractive future microbial chassis for producing carbon-neutral biofuels via synthetic biology and metabolic engineering, building upon our work using WT TIE-1 for bioplastic production^27^.

## Results

### Deleting an electron-consuming pathway enhances *n*-butanol production

We measured *n*-butanol production by WT with 3-gene cassette (WT+3), WT with 5-gene cassette (WT+5), and TIE-1 mutants with either 3-gene (Nif+3, Gly+3, Phb+3) or 5-gene cassette (Nif+5, Gly+5), under several photoheterotrophic and photoautotrophic conditions (substrate combinations, incubation time and final optical density listed in Supplementary Table 1-1, 1-2 and 1-3) to identify the most productive strains and conditions.

For photoheterotrophic conditions, we chose acetate (Ac) or 3-hydroxybutyrate (3Hy) as carbon and electron source because both substrates enter the *n*-butanol biosynthesis pathway directly as their CoA derivatives (Supplementary Fig. 1)^40^. For photoautotrophic conditions, we used either H_2_ or Fe(II) as an electron donor. CO_2_ was supplied in all conditions to maintain the pH of the medium and for redox balance in the cell. We provided N_2_ or ammonia (NH_4_^+^) as the nitrogen source.

We found that depending on the carbon and electron source used, the same construct produced variable amounts of *n*-butanol. *n*-Butanol production was the highest in the presence of 3Hy, followed by H_2_, Ac, and Fe(II) (Fig. 2, Supplementary Table 2-1 and 2-2). We found that Nif+5 is the most efficient *n*-butanol producer with highest production of 4.98 ± 0.87 mg/L under the photoheterotrophic conditions with NH_4_^+^ (Fig. 2b, Supplementary Table 2-1). The same construct, however, produced ~10-fold lower *n*-butanol when incubated with Fe(II) (0.55 ± 0.03 mg/L) (Fig. 2d, Supplementary Table 2-1 and 2-2).

**Figure 2.**
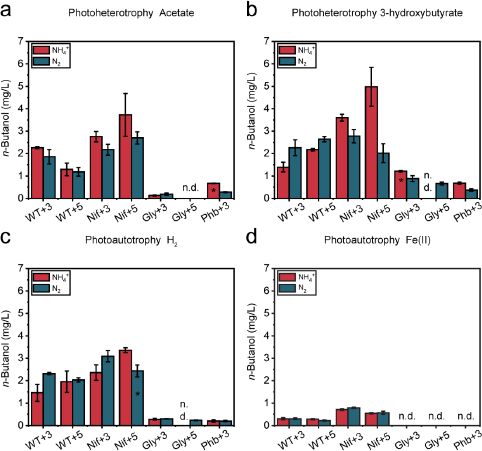
The nitrogenase double mutant (Nif) produced the highest amount of *n*-butanol in the presence of 3-hydroxybutyrate. The concentration of *n*-butanol in mg/L when TIE-1 was cultured with ammonium (NH_4_^+^, red) or dinitrogen gas (N2, blue) and (**a**) acetate (photoheterotrophy) (**b**) 3-hydroxybutyrate (photoheterotrophy) (**c**) hydrogen (H_2_) (photoautotrophy) and (**d**) ferrous iron (Fe(II)) (photoautotrophy). CO_2_ was present in all conditions. Data are means ± s.e.m. (standard error of the mean) of three biological replicates (bars that only have two biological replicates are indicated by ‘*’) and three technical replicates. WT+3: wild type with 3-gene cassette; WT+5: wild type with 5-gene cassette; Nif+3: nitrogenase knockout t with 3-gene cassette; Nif+5: nitrogenase knockout with 5-gene cassette; Gly+3: glycogen synthase knockout with 3-gene cassette; Gly+5: glycogen synthase knockout with 5-gene cassette; Phb+3: hydroxybutyrate polymerase knockout with 3-gene cassette, n.d. (non-detectable).

Compared to WT+3/WT+5, Nif+3/Nif+5 produced similar or more *n*-butanol depending on the substrates, whereas Gly+3/Gly+5 and Phb+3 produced less *n*-butanol from all substrates (Fig. 2, Supplementary Table 2-3). The presence of NH_4_^+^ in the media has been reported to repress the nitrogenase genes^41^. Therefore, we speculated that in its presence, WT+3/WT+5, and Nif+3/Nif+5 would produce similar amounts of *n*-butanol. Surprisingly, in most cases, Nif+3/Nif+5 produced a higher amount of *n*-butanol than the WT+3/WT+5, even in the presence of NH_4_^+^ (Fig. 2, Supplementary Table 2-3). Overall, we observe that deleting an electron-consuming pathway (Nif) is beneficial, whereas deleting an acetyl-CoA-consuming pathway (Gly and Phb) is detrimental to *n*-butanol production.

No *n*-butanol was detected from WT (no cassette added) using 3Hy as a carbon source. To ensure the *n*-butanol production is not toxic to TIE-1^7^, we performed a toxicity assay. The lowest inhibitory concentration of *n*-butanol for TIE-1 is 4050 mg/L (Supplementary Table 3-1), which is much higher than the highest *n*-butanol production (4.98 ± 0.87 mg/L). Hence, the *n*-butanol produced during our study does not limit the growth of TIE-1.

### Deleting acetyl-CoA-consuming pathways diverts carbon to acetone production

Acetone is a major byproduct of *n*-butanol biosynthesis^40^. As shown in Fig. 1a and Supplementary Fig. 1, the accumulation of acetoacetyl-CoA, an intermediate in *n*-butanol biosynthesis, leads to acetone production^7,40^. We observed highest acetone production by Phb+3 (0.00 to 290.01 ± 47.51 mg/L) followed by Gly+3/Gly+5 (1.47 ± 0.08 mg/L to 192.84 ± 4.82 mg/L), WT+3/WT+5 (0.00 mg/L to 107.39 ± 3.74 mg/L), and Nif+3/Nif+5 (0.00 mg/L to 76.44 ± 1.12 mg/L) (Fig. 3 Supplementary Table 2-4). This acetone production trend is in the reverse order of *n*-butanol production, i.e., Nif+3/Nif+5 produced the highest, and the Phb+3 produced the lowest amount of *n*-butanol (Fig. 2, Supplementary Table 2-3). These results indicate that acetone biosynthesis likely competes for acetyl-CoA with *n*-butanol biosynthesis. Using either Phb or Gly, acetyl-CoA that would have otherwise been directed toward PHB or glycogen synthesis was diverted to acetone biosynthesis.

**Figure 3.**
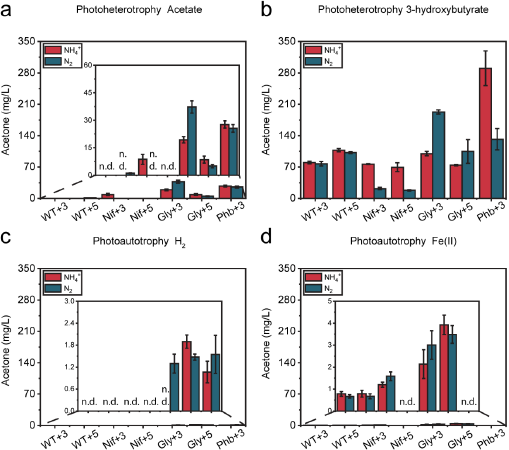
High *n*-butanol production correlates to low acetone production amongst TIE-1 mutants. The concentration of acetone in mg/L when TIE-1 was cultured with ammonium (NH_4_^+^, red) or dinitrogen gas (N2, blue) and (**a**) acetate (photoheterotrophy) (**b**) 3-hydroxybutyrate (photoheterotrophy) (**c**) hydrogen (H_2_) (photoautotrophy) and (**d**) ferrous iron (Fe(II)) (photoautotrophy). CO_2_ was present in all conditions. Data are means?± s.e.m. (standard error of the mean) of three biological replicates and three technical replicates. WT+3: wild type with 3-gene cassette; WT+5: wild type with 5-gene cassette; Nif+3: nitrogenase knockout t with 3-gene cassette; Nif+5: nitrogenase knockout with 5-gene cassette; Gly+3: glycogen synthase knockout with 3-gene cassette; Gly+5: glycogen synthase knockout with 5-gene cassette; Phb+3: hydroxybutyrate polymerase knockout with 3-gene cassette, n.d. (non-detectable).

Compared to using Ac as a substrate, which produced 0.00 mg/L to 37.29 ± 3.40 mg/L of acetone, all constructs produced ~10-100-fold more acetone when supplied with 3Hy (18.15 ± 1.41 to 290.10 ± 38.80 mg/L (Fig. 3a, 3b, Supplementary Table 2-5, 2-6). However, when the same strain was used, the acetone production under photoautotrophic conditions was lowered by ~25-125-fold compared to photoheterotrophic conditions with only 0.00 to 3.92 ± 0.44 mg/L (Fig. 3c, 3d) (Supplementary Table 2-5, 2-6). These results indicate that under photoheterotrophic conditions, particularly with 3Hy, TIE-1 accumulates more acetyl-CoA, which is eventually converted into acetone. The high acetone production suggests that acetyl-CoA is not limiting for *n*-butanol production. We also tested acetone toxicity in TIE-1 and found that the amount of acetone produced does not limit TIE-1’s growth (Fig. 3, Supplementary Table 3-2).

### More reducing equivalents enhance carbon conversion efficiency (CCE) to *n*-butanol

To further identify the most efficient strain and substrate for *n*-butanol production with respect to carbon, we determined carbon consumption (Supplementary Fig. 2) and CCE for each construct under all conditions.

#### Carbon consumption

We have recently shown that TIE-1 can fix CO_2_ during photoheterotrophic growth^30,41^. Therefore, we also calculated CO_2_ consumption and generation by all constructs. Under photoheterotrophy, all TIE-1 constructs consumed more (or generated less, represented by smaller negative value) CO_2_ with 3Hy (up to -114.23 ± 4.52 to 78.67 ± 15.86 μmol) than Ac (up to -243.67 ± 5.79 to 53.79 ± 9.77 μmol) (Fig. 4a, 4b, Supplementary Table 2-7, 2-8). With either 3Hy or Ac, Nif+3/Nif+5 consumed more CO_2_ (or generated less) CO_2_ generation (−50.53 ± 8.01 to 78.67± 15.86 μmol) (Fig. 4a, 4b, Supplementary Table 2-9). These results are consistent with a previous study where the use of a more reduced substrate (such as 3Hy) resulted in more carbon consumption than the use of a more oxidized substrate (such as acetate) for redox balance^41^.

**Figure 4.**
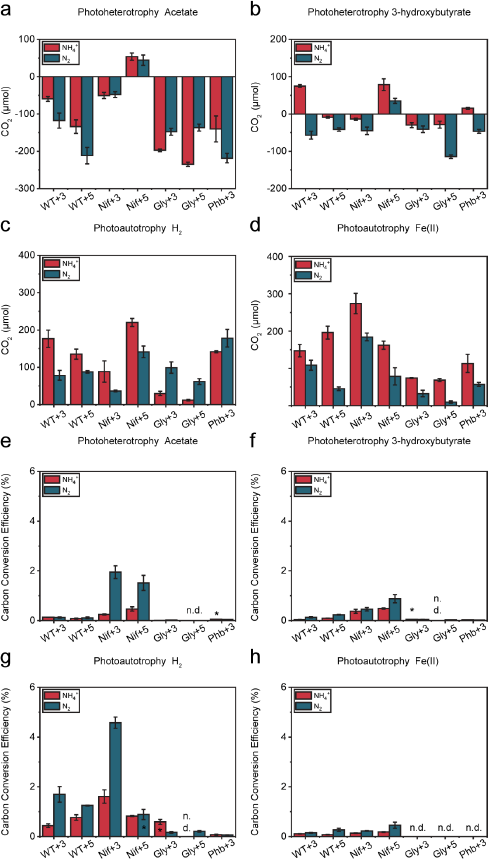
The nitrogenase double mutant (Nif) converts carbon to *n*-butanol more efficiently. **(a)-(d):** The CO_2_ consumption (positive value)/production (negative value). (**e**)-(**h**): The percentage of carbon converted to *n*-butanol. (**a**) and (**e**): acetate (photoheterotrophy) (**b**) and (**f**): 3-hydroxybutyrate (photoheterotrophy) (**c**) and (**g**): hydrogen (H_2_) (photoautotrophy) and (**d**) and (**h**): ferrous iron (Fe(II)) (photoautotrophy). CO_2_ was present in all conditions. CO_2_ was present in all conditions. Data are means± s.e.m. (standard error of the mean) of three biological replicates (bars that only have two biological replicates are indicated by ‘*’) and three technical replicates. WT+3: wild type with 3-gene cassette; WT+5: wild type with 5-gene cassette; Nif+3: nitrogenase knockout with 3-gene cassette; Nif+5: nitrogenase knockout with 5-gene cassette; Gly+3: glycogen synthase knockout with 3-gene cassette; Gly+5: glycogen synthase knockout with 5-gene cassette; Phb+3: hydroxybutyrate polymerase knockout with 3-gene cassette, n.d. (non-detectable).

Similarly, under photoautotrophic conditions, Nif+3/Nif+5 consumed the highest amount CO_2_ (36.41 ± 2.17 to 273.76 ± 27.25 μmol), except for Nif+3 incubated with H_2_ and NH_4_^+^ (Fig. 4c, 4d, Supplementary Table 2-9). This observation is likely due to the higher CO_2_-fixation required to achieve redox balance in the absence of N_2_-fixation. Gly+3/Gly+5 consumed the lowest amount of CO_2_ ranging from -234.67 ± 5.79 to 99.04 ± 15.32 μmol (Fig. 4c, 4d, Supplementary Table 2-9). Glycogen mutants have been reported to fix less CO_2_ compared to WT in cyanobacteria^36^. This observation corroborates our result that Gly+3/Gly+5 produces low *n*-butanol under photoautotrophic conditions (Fig. 2c, 2d, Supplementary Table 2-3).

#### CCE

Similar to the trend for *n*-butanol production (Fig. 2), Nif+3/Nif+5 showed the highest CCE (0.12 ± 0.03 to 4.58 ± 0.23%), followed by WT+3/WT+5 (0.03 ± 0.01 to 1.70 ± 0.32%), Gly+3/Gly+5 (0.00 to 0.59 ± 0.10%), and Phb+3 (0.00 to 0.16 ± 0.04%) (Fig. 4, e-h Supplementary Table 2-10). These results suggest that excess reducing equivalents enhanced *n*-butanol production and facilitated CCE to *n*-butanol. In contrast, lack of the PHB or glycogen biosynthesis decreased overall CCE to *n*-butanol. We found that all strains had the highest CCE when incubated with H_2_ (0.00 to 4.58 ± 0.23%), except for Phb (Fig. 4g, Supplementary Table 2-11, 2-12), which was unable to produce *n*-butanol using any substrate (Fig. 2c). This high CCE in the presence of H_2_ (Fig. 4g, Supplementary Table 2-11, 2-12) could be due to low acetone production (Fig. 3c, Supplementary Table 2-5, 2-6) under this condition.

Higher CCE (1- to 7-fold) was observed when Nif+3/Nif+5 was supplied with N_2_ compared to NH_4_^+^. For example, in the presence of NH_4_^+^, Nif+3/Nif+5 showed CCE of 0.14 ± 0.01 to 1.61 ± 0.27%, which increased to 0.23 ± 0.01 to 4.58 ± 0.23% when N_2_ was provided (Fig. 4 e-h, Supplementary Table 2-13). In summary, excess reducing equivalents in the Nif mutant leads to higher CCE towards *n*-butanol by TIE-1.

### More reducing equivalents enhance electron conversion efficiency to *n*-butanol

To further identify the most productive strain and substrate toward *n*-butanol production with respect to electron availability, we calculated each construct’s electron conversion efficiency (electron donor consumption data shown in Supplementary Fig. 2). We found that photoautotrophic conditions are more favorable for higher electron conversion efficiency than the photoheterotrophic conditions. With an electron conversion efficiency of 0.00 to 12.47 ± 1.37 %, Fe(II) was the most efficient condition followed by H_2_ (0.00 to 0.59 ± 0.14%), Ac (0.00 to 0.49 ± 0.06 %), and 3Hy (0.00 to 0.07 ± 0.01%) (Fig. 5, Supplementary Table 2-14, 2-15). The highest electron conversion efficiency was observed in the presence of Fe(II)^27^.

**Figure 5.**
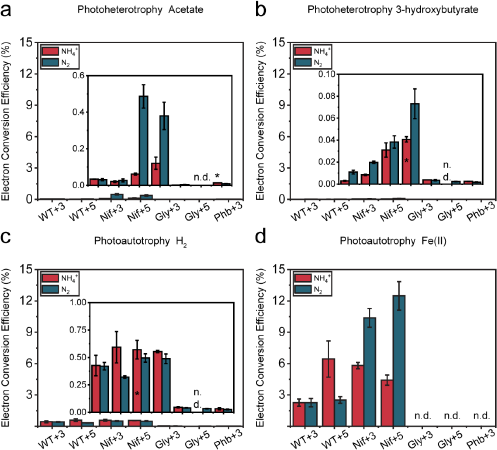
The nitrogenase double mutant (Nif) converts electron to *n*-butanol more efficiently. The electron conversion efficiency (%) when TIE-1 was cultured with ammonium (NH_4_^+^, red) or dinitrogen gas (N2, blue) and (**a**) acetate (photoheterotrophy) (**b**) (3-hydroxybutyrate (photoheterotrophy) (**c**) hydrogen (H_2_) (photoautotrophy); and (**d**) ferrous iron (Fe(II)) (photoautotrophy). CO_2_ was present in all conditions. Data are means?± s.e.m. (standard error of the mean) of three biological replicates (bars that only have two biological replicates are indicated by ‘*’) and three technical replicates. WT+3: wild type with 3-gene cassette; WT+5: wild type with 5-gene cassette; Nif+3: nitrogenase knockout t with 3-gene cassette; Nif+5: nitrogenase knockout with 5-gene cassette; Gly+3: glycogen synthase knockout with 3-gene cassette; Gly+5: glycogen synthase knockout with 5-gene cassette; Phb+3: hydroxybutyrate polymerase knockout with 3-gene cassette, n.d. (non-detectable).

Using the same carbon and electron source, the highest electron conversion efficiency was achieved by Nif+3/Nif+5 (0.04 ± 0.01 to 12.47 ± 1.37 %), followed by WT+3/WT+5 (0.00 to 6.45 ± 1.73%), Gly+3/Gly+5 (0.00 to 0.05 ± 0.00%), and Phb+3 (0.00 to 0.03 ± 0.01%, Fig. 5, Supplementary Table 2-16). In summary, availability of reducing equivalents due to deletion of an electron consuming pathway (Nif) leads to higher electron conversion efficiency for *n*-butanol biosynthesis in TIE-1.

### *n*-Butanol bioproduction can be achieved with light, electricity, and CO_2_

We have recently demonstrated that the photoelectroautotrophic growth of TIE-1 leads to a highly reduced intracellular environment compared to other growth conditions^30^. Our data show that excess reducing equivalents enhance *n*-butanol production in TIE-1. We further investigated *n-*butanol production by TIE-1 under photoelectroautotrophy using a three-electrode sealed BEC (Fig. 6a). We used Nif+5 in all BEC experiments as it was the most efficient *n*-butanol producer under most of the tested conditions (Fig. 2c, 4c, 5g, Supplementary Table 2-3, 2-10, 2-16).

**Figure 6.**
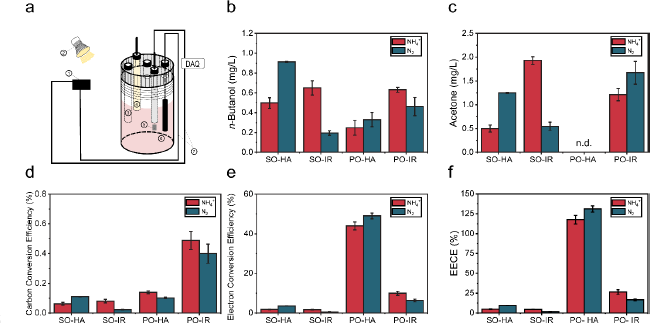
*n*-Butanol production, acetone production carbon conversion efficiency, electron conversion efficiency and electrical energy conversion efficiency (EECE) by the nitrogenase double mutant with the 5-gene cassette under photoelectroautotrophy. Under photoelectroautotrophic conditions, TIE-1 gains electrons from a poised electrode, using light as an energy source and carbon dioxide as a carbon source. For all the platforms, either ammonium (NH_4_^+^) or dinitrogen gas (N2) was supplied. (**a**) Schematic set up of BEC platform. platform set up: 1-electricity source 2-light source, 3-Purge inlet, 4-Reference electrode (Ag/AgCl in 3M KCl), 5-Counter electrode (Pt foil, 5 cm^2^), 6-Working electrode (Graphite rod, 3.2 cm^2^), DAQ-Data acquisition); (**b**) *n*-butanol production; (**c**) acetone production; (**d**) carbon conversion efficiency; (**e**) electron conversion efficiency; (**f**) Electrical Energy conversion efficiency EECE (%). PO-potentiostat, IR-infrared light, HA: halogen light, SO: solar panel. Data are means?± s.e.m. (standard error of the mean) of two biological replicates and three technical replicates.

We created four distinct biofuel production BEC platforms by combining two different electricity sources (grid-powered potentiostat or a solar panel) with two light sources (infrared or halogen light). The potentiostat approach represents conventional electrical sources, while the solar panel approach allows us to leverage renewably generated electricity. Infrared light is only a small portion of the solar spectrum that specifically excites the photosystem of TIE-1^26^. Halogen light mimics natural sunlight that represents the solar spectrum^42,43^. So, it can excite the photosystem of TIE-1 and support electricity generation by a solar panel, simultaneously. *BEC platform 1* used solar panel generated electrons and halogen light; *BEC platform 2* used solar panel generated electrons and infrared light; *BEC platform 3* used potentiostat and halogen light; *BEC platform 4* used potentiostat and infrared light. Either N_2_ or NH_4_^+^ was supplied as the nitrogen source. Supplementary Table 1-4 lists detailed platform setups. We measured *n*-butanol production, acetone production, and calculated CCE and electron conversion efficiency for each platform. We also calculated the electrical energy conversion efficiency (EECE) by dividing the combustion heat of the produced *n*-butanol by the electrical energy input.

The highest (0.91 ± 0.07 mg/L) and the lowest (0.19 ± 0.02 mg/L) *n*-butanol production was achieved when N_2_ was supplied as a nitrogen source in *BEC platform 1* and *BEC platform 2*, respectively (Fig. 6b, Supplementary Table 2-17, 2-18). The BEC platforms powered by solar panels showed 3-8 folds higher CO_2_ consumption (Supplementary Fig. 3a, Supplementary Table 2-17) and 5-40 folds higher electron uptake (Supplementary Fig. 3b, Supplementary Table 2-17)^30,41^ compared to the BEC platforms powered by the grid-powered potentiostat. Similar to the other autotrophic conditions (Fig. 3c, 3d), little or no acetone was produced (Fig. 6c) from the BEC platforms. BEC platform 4 achieved highest CCE (0.49 ± 0.06%, Fig. 6d, Supplementary Table 2-17, 2-18) compared to the other three BEC platforms. Although BEC platforms powered by grid powered potentiostat achieved much lower electron uptake (Supplementary Fig. 3b), they reached a much higher electron conversion efficiency (6-25 folds) than the platforms powered by a solar panel (Fig. 6e, Supplementary Table 2-17). This difference might be due to higher electrical loss associated with the solar panel.

BEC platforms under halogen light achieved higher electron conversion efficiency (4 to 8-fold, except using solar panel incubated with NH_4_^+^, Fig. 6e, Supplementary Table 2-18) compared to the BEC platforms using infrared light. However, the BEC platforms illuminated by halogen light (platforms 1 and 3) had much lower (20-90%) electron uptake, particularly when using solar panel as electricity source (Supplementary Fig. 3b Supplementary Table 2-18). To ensure that a lower number of attached cells did not reduce electron uptake from the platforms using halogen light, we performed live-dead viability assay. We observed that the percentage and number of live cells attached to the electrodes were similar in all the BEC platforms (40%-50%) (Supplementary Fig. 3c, and 3d, Supplementary Table 2-18). These results indicate that halogen light is not the ideal light source for TIE-1 with respect to electron uptake.

We further compared the EECE between the two electricity sources. We found that the BEC platforms powered by solar panel show lower EECE (1.62 ± 0.20 to 9.55 ± 0.34%) than the BEC platforms using a potentiostat (16.62 ± 1.01 to 131.13 ± 3.97%) when the same nitrogen source (either N_2_ or NH_4_) was supplied (Fig. 6f, Supplementary Table 2-17). With respect to the light source, platforms using halogen light resulted in higher EECE (4.80 ± 0.38 to 131.14 ± 3.97%) than platforms using infrared light (1.62 ± 0.20 to 26.52 ± 2.87%) when the same nitrogen source was supplied (6f, Supplementary Table 2-18). Halogen light represents the solar spectrum and several wavelengths from this light source can be absorbed by TIE-1 via the light-harvesting complexes and eventually, the photosystem^30,44^. This would lead to higher ATP synthesis via cyclic photosynthesis by TIE-1^30^ perhaps explaining the greater than 100% EECE.

In summary, *BEC platform 1* showed higher *n*-butanol production, *BEC platform 4* showed the highest CCE, and *BEC platform 3* showed the highest electron conversion efficiency and EECE. Although *BEC platform 1* resulted in moderate conversion efficiencies, the highest *n*-butanol production (up to 5-fold) with the use of sustainable resources (electricity from solar panel and energy from halogen light), make this platform the most promising for further development as a sustainable and carbon-neutral process for *n*-butanol production.

## Discussion

In recent years, *n*-butanol has been proposed as a superior biofuel due to its higher energy content, lower volatility, and reduced hydrophilicity^4^. Here we produced *n*-butanol by introducing an artificial *n*-butanol biosynthesis pathway^34^ into an anoxygenic photoautotroph *Rhodopseudomonas palustris* TIE-1^26^. Using metabolic engineering and novel hybrid bioelectrochemical platforms, we show that TIE-1 can produce *n*-butanol using different carbon sources (organic acids, CO_2_), electron sources [H_2_, Fe(II), a poised electrode], and nitrogen sources (NH_4_^+^, N_2_). TIE-1’s ability to produce *n*-butanol under photoelectroautotrophy using light, electricity, and CO_2_ can serve as a stepping-stone for sustainable solar fuel production.

After introducing a codon-optimized *n*-butanol biosynthesis pathway in TIE-1 and its mutants (Nif, Gly, and Phb), we determined *n*-butanol production, acetone production, CCE, and electron conversion efficiency of these constructs under both photoheterotrophic and photoautotrophic conditions. Mutants lacking the nitrogen-fixing pathway (Nif+3/Nif+5) (known to affect redox balance in the cell by consuming reducing equivalents^37,39,45^) exhibited a more reduced intracellular environment (indicated by higher CO_2_ fixation)^41^ and produced more *n*-butanol compared to WT+3/WT+5. In contrast, deleting acetyl-CoA-consuming pathways (Gly+3/Gly+5 and Phb+3) led to lower *n*-butanol production. These results show that higher reducing equivalent rather than increased acetyl-CoA availability enhances *n*-butanol production by TIE-1. These results also agree with previous works where redox balance or reducing equivalent availability plays a vital role in *n*-butanol production^33,46^. A closely related strain *R. palustris* CGA009 has been shown to produce *n*-butanol when its biosynthesis was the obligate route for maintaining redox balance during photoheterotrophic growth on *n*-butyrate^33^. Similarly, in *E. coli, n*-butanol production increased when its biosynthesis acted as an electron-sink to rescue cells from redox imbalance^46^.

We expected that the presence of NH_4_^+^ would inhibit the expression of nitrogenase, so nitrogen fixation would not occur. Thus, nitrogenase would not produce the byproduct H_2_^45,47,48^. However, we observed that WT+3/WT+5 and Gly+3/Gly+5 produced H_2_ (likely via nitrogenase) despite the presence of NH_4_^+^ (Supplementary Fig. 2). This was in contrast to the Nif+3/Nif+5, which did not produce H_2_ under any condition, confirming that the observed H_2_ production in the WT and Gly strains is due to nitrogenase activity. The production of H_2_ by nitrogenase is well known in CGA009^38,39,41^. This unexpected nitrogenase activity could have been initiated by the lower NH_4_^+^ concentrations toward the end of the experiment, which might lead to the induction of nitrogenase gene expression^47,49^. Because H_2_ production via nitrogenase even in the presence of NH_4_^+^ consumes reducing equivalents, this explains why the Nif strains produce more *n*-butanol compared to the WT and Gly strains.

We also observed that by feeding *n*-butanol biosynthesis pathway intermediates as a carbon source, such as 3Hy, TIE-1 produced more *n*-butanol (Fig. 2). However, despite high *n*-butanol production with 3Hy, TIE-1 showed low CCE and low electron conversion efficiency, possibly due to higher acetone production (Fig. 3). This high acetone production is likely due to the accumulation of acetoacetyl-CoA, converted from 3Hy through 3-hydroxybutyryl-CoA (Supplementary Fig. 1)^50^. This along with 3Hy being an expensive feedstock compared to CO_2_ for bioproduction^51,52^ makes it an unsuitable substrate for economical *n*-butanol production.

In general, we achieved higher *n*-butanol production, CCE, and electron conversion efficiency when acetone production was lower. These results agree with the previous studies where an increase in *n*-butanol production accompanies a decrease in acetone production^53,54^. Although using highly reduced substrates, such as glycerol, can increase the ratio of *n*-butanol to acetone, a significant amount of acetone is always detected while using the *n*-butanol biosynthesis pathway from *C. acetobutylicum*^53,54^. Our study addressed this issue by using slow or non-growing cells that produced *n*-butanol without the production of acetone.

BEC platforms powered by the potentiostat resulted in higher EECE and electron conversion efficiency. This difference might be due to either electrical or optical losses associated with the solar panel during photoelectron generation^55,56^. The electrical loss could be due to the limited energy efficiency of the solar panel, which is determined by the diode characteristics and series resistances in the solar panel^55,56^. And optical loss can be in the form of poor light absorbance or light reflection from the solar cell surfaces or material defects^57,58^. We found that the platforms with halogen as the light source have higher EECE (~8 fold) regardless of the electricity source.

To contextualize our results, we compared EECE, E_appl_, and *n*-butanol production, and CCE with the previous related studies.

### EECE

Using solar panel generated electricity TIE-1 achieved an EECE of up to 9.54%, which increased by over 13-fold (up to 131.13%) when we used grid-based electricity (Fig. 6f). In a previous study using a hybrid water-splitting system, *R. eutropha* achieved an EECE of 16% towards C_4_+C_5_ alcohol using grid-based electricity^19^. These data suggest that TIE-1 can achieve higher EECE.

### E_appl_ and power requirement

TIE-1 can gain electrons directly from an electrode, which requires lower E_appl_ for photoautotrophic growth and *n*-butanol biosynthesis (E_appl_ = 0.1 − 0.5 V). In contrast, the hybrid water splitting system used to synthesize C_3_-C_5_ alcohol or PHB by *R. eutropha* used an E_appl_ of 2.0 V. Similarly, *n*-butanol synthesis by *Clostridium* sp. using MES used an E_appl_ of 0.8 V12,19. Assuming that all the reactors use 1 mA of current, the power would be 5 x 10^-4^ W for *n*-butanol bioproduction by TIE-1. In contrast, *R. eutropha* would require 2 x 10^-3^ W for water-splitting, and *Clostridium* sp. would require 8 x 10^-4^ W. Therefore, TIE-1 uses four times less power than *R. eutropha* and 1.6 times less power than *Clostridium* sp. This implies that even low-efficiency solar panel-based platforms^57^, and low sunlight conditions can more be easily used for bioproduction using organisms like TIE-1^59,60^.

### n-Butanol production

Under photoelectroautotrophy, TIE-1 produced 0.91 ± 0.07 mg/L of *n*-butanol in 10 days (Fig. 6b). *Clostridium* sp. produced 135 mg/L *n*-butanol in 35 days^12^. Compared to *R. eutropha* and *Clostridium* sp., our platform produced lower *n*-butanol. Under photoautotrophic conditions, TIE-1 produced a maximum of 3.09 ± 0.25 mg/L of *n*-butanol in batch culture (Fig. 2c). Initial studies in cyanobacteria resulted in 2.2 mg/L^14^. Recently, using a modular engineering method, cyanobacteria can produce 4.8 g/L of *n*-butanol^17^, which is 2000 times higher than the initial production. With intensive future engineering efforts, we anticipate that TIE-1 can also demonstrate higher *n*-butanol production.

### CCE

To the best of our knowledge, no autotrophic *n*-butanol production study has reported CO_2_ consumption^14–17,35,36^. Thus, here we compared TIE-1’s CCE with that reported for heterotrophic *n*-butanol production. Although most heterotrophic growth media use yeast extract (an undefined carbon source), for simplicity, the CCE calculations considered glucose as the only carbon source. We calculated CCE using the total amount of added carbon in these studies^9,61^. Considering the additional contribution of yeast extract would lower the CCE further. The early trials in *E. coli* and *S. cerevisiae* reached carbon conversion efficiencies of 0.11 % and 0.02%. With intensive metabolic engineering, the CCE reached 45.92% (*E. coli*) and 11.52% (*S. cerevisiae*) (calculated from the reported g/g yield)^34,62^. In this initial study here, we show that TIE-1 shows CCE (mol/mol) of 4.58 ± 0.21% and 1.95 ± 0.26% under photoautotrophic and photoheterotrophic conditions, respectively (Fig. 4). This is 20 and 200 times higher than that of initial studies in *E. coli* and *S. cerevisiae*. Photoautotrophic bioproduction is superior due to the low cost of CO_2_ compared to heterotrophic substrates^52^. Thus, developing TIE-1 further via metabolic engineering, synthetic biology, and bioprocess engineering will make it an economically viable bioproduction platform.

In summary, TIE-1 can achieve high electrical energy conversion efficiency, and CCE with lower power input, while producing an amount of *n*-butanol comparable to the initial studies in established bioproduction chassis organisms like *E. coli*, and *S. cerevisiae*. This study represents the initial effort of producing carbon-neutral fuels using TIE-1. Countless modifications could be made to improve the *n*-butanol titer. For example, we observed an increased expression of genes in the *n*-butanol biosynthesis pathway from Nif+5 incubated with 3Hy (the strain and condition that resulted in the highest *n*-butanol production) (Supplementary Fig. 4). Therefore, increasing gene expression by driving each gene in the *n*-butanol biosynthesis pathway with its own promoter could increase *n*-butanol production. Also, since reducing equivalent availability seems to be a bottleneck for *n*-butanol production, deleting more pathways that consume electrons could increase *n*-butanol production. Also, increasing intracellular iron could lead to higher cytochrome production, which would increase electron uptake^63^. Furthermore, creating a BEC platform with built-in solar conversion to electricity capability could reduce electrical energy loss. Finally, higher electron uptake, which should be beneficial for *n*-butanol synthesis, could be achieved by using nanoparticle modified electrodes^31,64,65^. Taken together, TIE-1 offers a sustainable route for carbon-neutral *n*-butanol biosynthesis and other value-added products. As CO_2_ concentrations are rising in the atmosphere, such bioproduction strategies need immediate attention and support.

## Materials and Methods

### Bacterial strains, media, and growth conditions

All strains used in this study are listed in Supplementary Table 1-5 *E. coli* strains were grown in lysogeny broth (LB; pH 7.0) at 37°C. For aerobic growth, *Rhodopseudomonas palustris* TIE-1 was grown at 30 °C in YP medium (3 g/L yeast extract, 3 g/L peptone) supplemented with 10 mM MOPS [3-N (morpholino) propanesulphonic acid] (pH 7.0) and 10 mM succinate (YPSMOPS) in the dark. For growth on a solid medium, YPSMOPS or LB was supplemented with 15 g/L agar. For anaerobic phototrophic growth, TIE-1 was grown in anoxic bicarbonate buffered freshwater (FW) medium^27^. All FW medium was prepared under a flow of 34.5 kPa N_2_ + CO_2_ (80%, 20%) and dispensed into sterile anaerobic Balch tubes. The cultures were incubated at 30°C in an environmental chamber fitted with an infrared LED (880 nm). For photoheterotrophic growth, the FW medium was supplemented with 50 mM MOPS at pH 7.0 and sodium 3-hydroxybutyrate or sodium acetate at pH 7.0, to a final concentration of 50 mM. For photoautotrophic growth on iron, anoxic sterile stocks of FeCl_2_ and nitrilotriacetic acid (NTA) were added to reach final concentrations of 5 mM and 10 mM, respectively. For photoautotrophic growth on H_2_, TIE-1 was grown in FW medium at pH 7.0 and 12 psi of 80% H_2_/20% CO_2_^27^. For all carbon and electron sources, either ammonium chloride (5.61 mM) or dinitrogen gas (8 psi) was supplied as nitrogen source^27^. All sample manipulations were performed inside an anaerobic chamber with a mixed gas environment of 5% H_2_/75% N_2_/20% CO_2_ When needed, 400 μg/mL kanamycin was added for TIE-1, and 50 μg/mL kanamycin was added for *E. coli*.

### *R. palustris* TIE-1 deletion mutant construction

We constructed three mutants, two of which were double mutants using the method described in a previous study^30^. Respectively, Glycogen synthase knockout was created by deleting Rpal_0386, nitrogenase knockout was created by deleting Rpal_1624, and Rpal_5113, and hydroxybutyrate polymerase knockout was created by deleting and Rpal_2780 and Rpal_4722 were deleted resulting hydroxybutyrate polymerase knockout. Briefly, the 1 kb upstream and 1 kb downstream regions of the gene were PCR amplified from the *R. palustris* TIE-1 genome, then the two homology arms of the same gene were cloned into pJQ200KS plasmid. The resulting vector was then electroporated into *E. coli* and then conjugated to *R. palustris* TIE-1, using the mating strain *E. coli* S17-1/λ. After two sequential homologous recombination events, mutants were screened by PCR, as shown in Supplementary Fig. 5. The primers used for mutant construction and verification are listed in Supplementary Table 1-6 and 1-7.

### Plasmid construction

All plasmids used in this study are listed in Supplementary Table 1-8. There are five genes involved in the *n*-butanol biosynthesis: *phaJ, ter, adhE2, phaA*, and *phaB* (Fig. 1a). Among these five genes, TIE-1 has homologs of the first two (*phaA* and *phaB*). Hence, we designed two different cassettes, namely, a 3-gene cassette (3-gene), which has *phaJ, ter, adhE*2, and a 5-gene cassette (5-gene), which has the 3-gene plus a copy of the *phaA-phaB* operon from TIE-1. *phaJ, ter*, and *adhE2* sequences were obtained from published studies^34^. The *phaJ* gene, isolated from *Aeromonas caviae*, was chosen because it codes for an enzyme that has a higher specificity for its substrate^34,66^. The *ter* gene isolated from *Euflena gracilis* was selected because it is unable to catalyze the reverse oxidation of butyryl-CoA^34^. The *adhE*2 gene isolated from *C. acetobutylicum* chosen because the enzyme encodes for specifically catalyzes the reduction of the butyryl-CoA^34^. All three foreign genes (*phaJ, ter*, and *adhE2*) were codon-optimized by Integrated DNA Technology (IDT) for TIE-1. The cassette was synthesized as G-blocks by IDT, which we then stitched together by overlap extension and restriction cloning. The *phaJ-ter-adhE2* cassette was then inserted into plasmid pRhokS-2, resulting in pAB675. *PhaA* and *phaB* were amplified as an operon from the *R. palustris* TIE-1 genome. The *phaA-phaB* cassette was then cloned into pAB675 to obtain pAB744. Upon obtaining mutants and plasmids, either the 3-gene or the 5-gene was conjugated into WT TIE-1 or the mutants, using mating the strain *E. coli* S17-1/ λ. All conjugations were successful, except for the 5-gene into the Δ*phaC*1Δ*phaC*2. The primers used for cassette construction are listed in Supplementary Table 1-9. The primers used for cassette sequencing are listed in Supplementary Table 1-10.

### Substrate measurement

Substrate concentrations at the beginning (T_0_) and the end (T_f_) were measured to calculate carbon and electron conversion efficiency to *n*-butanol. The incubation time of each experiment can be found in Supplementary Table 1-2.

#### a) CO_2_ and H_2_ analysis by gas chromatography

CO_2_ and H_2_ were analyzed using a method described in a previous study^27^. Gas samples were analyzed using gas chromatography (Shimadzu BID 2010-plus, equipped with Rt^®^-Silica BOND PLOT Column, 30 m × 0.32 mm; Restek, USA) with helium as a carrier gas. To measure the CO_2_ content of the liquid phase, 1 mL of the cell-free liquid phase was added to 15 mL helium-flushed septum-capped glass vials (Exetainer, Labco, Houston) containing 1 mL 85% phosphoric acid. Then 40 μL of the resulting gas from the Balch tube was injected into the Shimazu GC-BID, using a Hamilton^TM^ gas-tight syringe. To measure the CO_2_ and H_2_ contents of the gas phase, either 40 μL of the gas phase was directly injected into the Shimadzu GC-BID, or 5 mL of the gas phase was injected into a 15 mL helium-flushed septum-capped glass vial (Exetainer, Labco, Houston), using a Hamilton^TM^ gas-tight syringe. Then 50 μL of the diluted gas sample was injected into the Shimazu GC-BID, using a Hamilton^TM^ gas-tight syringe. A standard curve was generated by the injection of 10 μL, 25 μL, and 50 μL of H_2_ + CO_2_ (80%, 20%). The total moles of CO_2_ in the reactors were calculated using the ideal gas law (PV=nRT)^67^.

#### b) Organic acid analysis by ion chromatography

The acetate and 3-hydroxybutyrate concentrations were measured as described previously^27^. Briefly, after 1:50 dilution, the acetate and 3-hydroxybutyrate concentrations at the staring and endpoint of culture for each sample were quantified using an Ion Chromatography Metrohm 881 Compact Pro with a Metrosep organic acid column (250 mm length). Eluent (0.5 mM H_2_SO_4_ with 15% acetone) was used at a flow rate of 0.4 mL min^−1^ with suppression (10 mM LiCl regenerant)^27^.

#### c) Ferrous iron [Fe(II)] analysis by ferrozine assay

The Fe(II) concentration was measured using ferrozine assay, as described in a previous study^27^. Briefly, 10 μL of culture was mixed with 90 μL 1M HCl in a 96-well plate inside the anaerobic chamber with 5% H_2_/75% N_2_/20% CO_2_ (Coy laboratory, Grass Lake). After the plate was removed from the anaerobic chamber, 100 μL of ferrozine (0.1% (w/v) ferrozine in 50% ammonium acetate) was added to the sample. Then the 96-well plate was covered with foil and incubated at room temperature for 10 minutes before the absorbance was measured at 562 nm. The absorbance was then converted to Fe(II) concentration based on a standard curve generated by measuring the absorbance from 0 mM, 1 mM, 2.5 mM and, 5 mM Fe(II).

### *In vivo* production of *n*-butanol

The plasmids with the *n*-butanol pathway were unstable when adapting the strain to the nitrogen-fixing or photoautotrophic conditions. To avoid this problem, a twice-washed heavy inoculum from YPSMOPS was used under all conditions. All strains were inoculated in 50 mL of YPSMOPS with kanamycin with a 1:50 dilution from a pre-grown culture. When the OD_660_ reached 0.6-0.8, the culture was inoculated into 300 mL of YPSMOPS with kanamycin. When the OD_660_ reached 0.8~1, 10 mL of culture was saved for a PCR check (Supplementary Fig. 6). The rest of the culture was washed twice with ammonium-free FW medium and resuspended using anoxic ammonium free FW medium inside the anaerobic chamber. Finally, the culture was inoculated into the medium containing different carbon sources and electron donors (acetate, 3-hydroxybutyrate, H_2_, Fe(II), or electrode) in either a sealed Balch tube (initial OD_660_ ~1) or a bioelectrochemical cell (initial OD_660_ ~0.7). The tubes and the reactors were sealed throughout the process, and samples were taken after the cultures reached the stationary phase (incubation time listed in Supplementary Table 4), using sterile syringes.

### Extraction and quantification of *n*-butanol and acetone

After the culture entered the late stationary phase, 1 mL of culture was removed from the culture tube using a syringe and centrifuged at 21,100X *g* for 3 minutes. The supernatant was then filtered using a syringe filter, and the filtrate or the standard was extracted with an equal volume of toluene (containing 8.1 mg/L iso-butanol as an internal standard) and mixed using a Digital Vortex Mixer (Fisher) for 5 minutes, followed by centrifugation at 21,100X *g* for 5 minutes. After centrifugation, 250 μL of the organic layer was added to an autosampler vial with an insert. The organic layer was then quantified with GC-MS (Shimazu GCMS-QP2010 Ultra), using the Rxi^®^-1ms column. The oven was held at 40 °C for 3 minutes, ramped to 165 °C at 20 °C/min, then held at 165 °C for 1 min. Samples were quantified relative to a standard curve for 0 mg/L, 0.2025 mg/L, 0.405 mg/L, 0.81 mg/L, 2.025 mg/L, 4.05 mg/L, and 8.1 mg/L of *n*-butanol and 0 mg/L, 0.784 mg/L, 3.92 mg/L, 7.84 mg/L, 39.2 mg/L, 78.4 mg/L, and 392 mg/L of acetone. An autosampler was used to reduce the variance of injection volumes.

### Bioelectrochemical platforms and growth conditions

A three-electrode sealed-type bioelectrochemical cell (BEC, C001 Seal Electrolytic Cell, Xi’an Yima Opto-electrical Technology Com., Ltd, China)^30,64^ containing 80 mL of FW medium was used for testing *n*-butanol production. The three electrodes were configured as a working electrode (a graphite rod, 3.2 cm^2^), a reference electrode (Ag/AgCl in 3.5M KCl), and a counter electrode (Pt foil, 5 cm^2^). FW medium (76 mL) was dispensed into sterile, sealed, three-electrode BECs, which were bubbled for 60 minutes with N_2_ + CO_2_ (80%/20%) to remove oxygen and pressurized to ~7 psi. Four BECs were operated simultaneously (*n*=3 biological replicates) with one no-cell control. All photoelectroautotrophic experiments were performed at 26 °C under continuous infrared light (880 nm) or halogen light. The electrical potential of 0.5 V (E_appl_=0.5 V) was constantly applied between the working electrode and counter electrode using a grid powered potentiostat (Interface 1000E, Gamry Multichannel potentiostat, USA) or solar panel (Uxcell 0.5V 100mA Poly Mini Solar Cell Panel Module) for 240 hrs. Electron uptake and current density were collected every 1 minute using the Gamry Echem Analyst^TM^ (Gamry Instruments, Warmister, PA) software package. At the end of the bioelectrochemical experiment, the samples were immediately collected from the BEC reactors. *n*-butanol, acetone, and substrates were measured as described above.

### Calculations of CCE, electron conversion efficiency, and electrical conversion efficiency

CCE, electron conversion efficiency, and electrical energy conversion efficiency (EECE) were calculated by dividing the total carbon/electrons/electrical-energy consumption by the final carbon/electrons/energy content in *n*-butanol, respectively.

To determine carbon consumption, acetate, 3-hydroxybutyrate, or CO_2_ consumption was calculated by subtracting the amount in the sample at the end of the experiment from the amount at the beginning of the experiment. Then all the carbon substrate consumptions were converted to moles of carbon, using Equation 1. The amount of carbon converted to *n*-butanol was calculated based on the *n*-butanol production, using Equation 2. The CCE was calculated using Equations 1, 2, and 3 below:

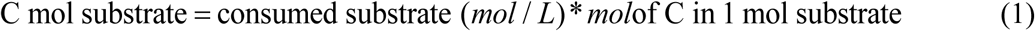

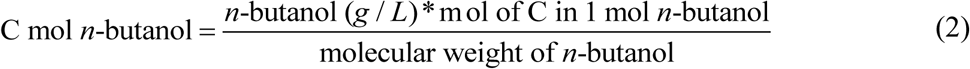

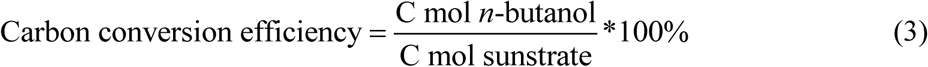

The theoretical total number of electrons available from each consumed electron donor was calculated as described below (Equation 4). The total available electrons from the complete oxidation of each organic acid were calculated with the assumption that the final oxidation product was CO_2_. The inorganic electron donors such as Fe(II) and H_2_ release 1 mole e^-^ and 2 moles e^-^ per mole, respectively. Electrons supplied for the photoelectroautotrophy condition were calculated directly from BEC based experiments wherein the total current uptake was integrated over the operational time. The total electron uptake was used to calculate the electron conversion efficiency to *n*-butanol because the electrode is the direct electron donor under this growth condition. The number of electrons required for *n*-butanol production was calculated from the oxidation state of the carbon in each carbon source and *n*-butanol. Supplementary Table 4 lists the specific oxidation state, and the number of electrons required per mole of *n*-butanol is listed for all studied sources and *n*-butanol.

To calculate total available electrons from each substrate, the amount of consumed substrate (in moles) was multiplied by the theoretical total available electrons per mole of the substrate when fully oxidized to CO_2_ (Equation 4). For photoelectroautotrophy, the total available electron was calculated based on data collected from a data acquisition system (DAQ, Picolog Datalogger). To obtain the electrons required for *n*-butanol production, the *n*-butanol production (in moles) was multiplied by the theoretical number of electrons required per mole (Equation 5). The conversion efficiency was calculated by dividing the moles of electrons required for *n*-butanol production by the theoretical total available electrons (Equation 6).

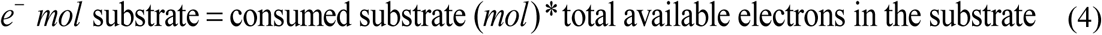

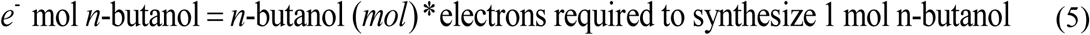

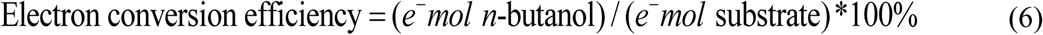

Calculation of the electrical energy conversion efficiency (EECE) to *n*-butanol was adapted from a previous study^19^. The EECE was calculated by equation 7. The charge supplied to the bioelectrochemical platforms was calculated from data collected by DAQ.

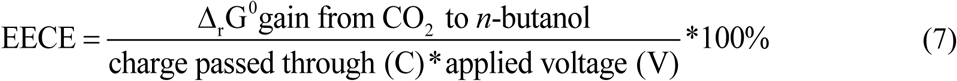

The Gibbs free energy gains (Δ_r_G^0^) for *n*-butanol was calculated similarly with a previous study^19^ by reaction 8 and equation 9^68^.

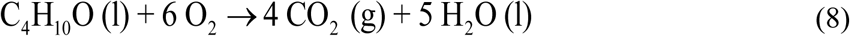

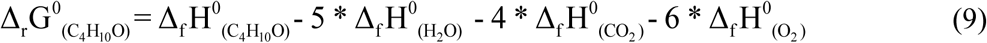

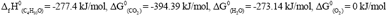

### RNA extraction, cDNA synthesis, and RT-qPCR

To extract RNA for cDNA synthesis and eventually perform RT-qPCR for analyzing the expression level of the individual genes, culture samples (2.5 ml to 15 ml depending on OD_660_) were taken at the late exponential (T_m_) or stationary phase (T_f_). Samples were immediately stabilized with an equal volume of RNAlater (Qiagen, USA). After incubation at room temperature for 10 min, samples were centrifuged at 21,100 *g for three minutes. After the supernatant was removed, the pellet was stored at -80°C before RNA extraction using the Qiagen RNeasy Mini kit (Qiagen, USA), following the manufacturer’s protocol. DNA was removed using a Turbo DNA-free Treatment and Removal Kit (Ambion, USA). DNA contamination was ruled out by PCR using the primers listed in Supplementary Table 1-11.

Purified RNA samples were then used for cDNA synthesis by an iScript^TM^ cDNA Synthesis Kit (Biorad, USA). The same mass of RNA was added to each cDNA synthesis reaction. The synthesized cDNA was used for RT-qPCR. RT-qPCR was performed using the Biorad CFX connect Real-Time System Model # Optics Module A with the following thermal cycling conditions: 95 °C for 3 min, then 30 three-step cycles of 95 °C for 3 seconds, 60 °C for 3 min, and 65 °C for 5 seconds, according to the manufacturer’s manual. The reaction buffer was iTaq SYBR Green Supermix with ROX (Bio-Rad). The primers used for RT-qPCR (listed in Supplementary Table 1-11) were designed using primer3 software (http://bioinfo.ut.ee/primer3/). The primer efficiencies were determined by performing RT-qPCR using different DNA template concentrations. The genes *clpX* and *recA*, which have been previously validated as internal standards, were used for the genome^29,30^. The gene code for kanamycin resistance was also used as an internal standard for the plasmid. After RT-qPCR, the data were analyzed using the C_T_ method. Fold changes, and standard deviations were calculated as described in a previous study^27^.

### Viability analysis of TIE-1 under photoelectroautotrophy

WT TIE-1 was inoculated into the bioelectrochemical reactors described above, with a starting OD of ~0.3. After 72 hours of incubation, the viability of the biofilm of the electrode was characterized by imaging the electrode stained with LIVE/DEAD^®^ (L7012, Life Technologies), and then the attached cells were quantified using NIS-Elements AR Analysis 5.11.01 64-bit software. Imaging of the electrode was performed as described in a previous study^30^. Prior to cutting a piece of the spent electrode, the electrode from the reactor was washed 3 times with 1X phosphate-buffered saline (PBS) to remove unattached cells. A piece of the spent electrode was then submerged in 1X PBS in a sterile microfuge tube. Prior to imaging, the electrode piece was immersed in LIVE/DEAD^®^ stain (10 μM SYTO9 and 60 μM propidium iodide) kit and incubated for 30 minutes in the dark. The electrode sample was then placed in a glass-bottom Petri dish (MatTek Corporation, Ashland, MA) containing enough PBS to submerge the sample. Further, it was imaged on a confocal microscope (Nikon A1 inverted confocal microscope), using 555 and 488 nm lasers and a 100X objective lens (Washington University in St. Louis Biology Department Imaging Facility). Electrode attached cells were quantified by Elements Analysis software using the protocol described below: Briefly, for each reactor, three images were processed. Z-stacks of each image were split into two channels (one for live cells, one for dead cells), the MaxIP was acquired for the combined z-stacks. After GaussLaplace, local contrast and smoothing, and thresholding, and Object Count was performed for each channel based on a defined radius (0.8μm~5μm). Then the percentage of live (or dead) cells was calculated by

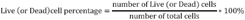

The viability of planktonic cells was determined by RT-qPCR of the essential genes (ATP synthase homologs: *atp*1, *atp*2) and the genes involved in photoelectroautotrophy (photosynthetic reaction center: *puf*L, ribulose-1,5-bisphosphate carboxylase/oxygenase: *ruBisCo*1, *ruBisCo*2, and pio operon: *pioA*) (Supplementary Fig. 3).

### Toxicity Study

WT TIE-1 with an empty vector (pRhokS-2) was used to test the tolerance of TIE-1 for acetone and *n*-butanol. To test the tolerance, 0%, 0.25%, 0.5%,1%, or 2% *n*-butanol (v/v), or 0%, 0.1%, 0.25%, 0.5%, 1%, or 2% acetone (v/v), was added to FW media with acetate (10 mM). Growth was monitored by recording OD_660_ over time.

### Statistics

All statistical analyses (two tails Student’s t-test) were performed with Python. *p*-value<0.05 was considered to be significant. All the experiments were carried out using biological triplicates and technical triplicates except for RT-qPCR for photoelectroautotrophy which was performed as technical duplicates.

## General

We thank the following members of the Washington University community: Marta Wegorzewska and James Ballard for their careful reading of the manuscript; Dianne Duncan for her help with confocal microscopy; and Dr. Joshua Blodgett, Dr. Michael Singh Guzman, Dinesh Gupta, and Yunci Qi for their helpful comments during the preparation of this manuscript.

## Author contributions

W.B.., A.B., and K.R. designed the research. W.B, T.O.R., and K.R, collected the data. W.B., T.O.R., and A.B. analyzed and interpreted the data. W.B., R.S., K.R. and A.B. wrote the manuscript. All authors reviewed, revised, and approved the final manuscript.

## Data and materials availability

All data in this study are available from the corresponding authors upon request.

