## Supplemental Materials for "Sustainable Production of the Biofuel *n*-Butanol by *Rhodopseudomonas palustris* TIE-1"

Bai, et al.

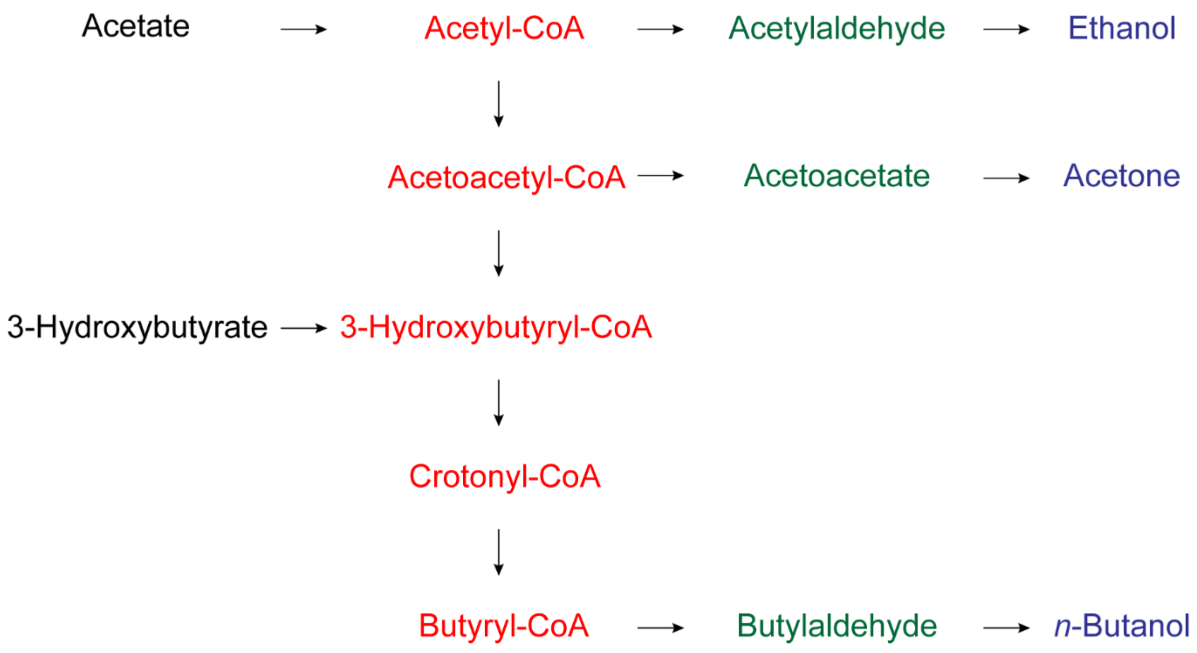


**Supplementary Figure 1. Substrates and by-products of *n*-butanol biosynthesis.** Black: substrates; Red: *n*-butanol synthesis intermediates; Green: in-cell by-products; Blue: secreted by-products.

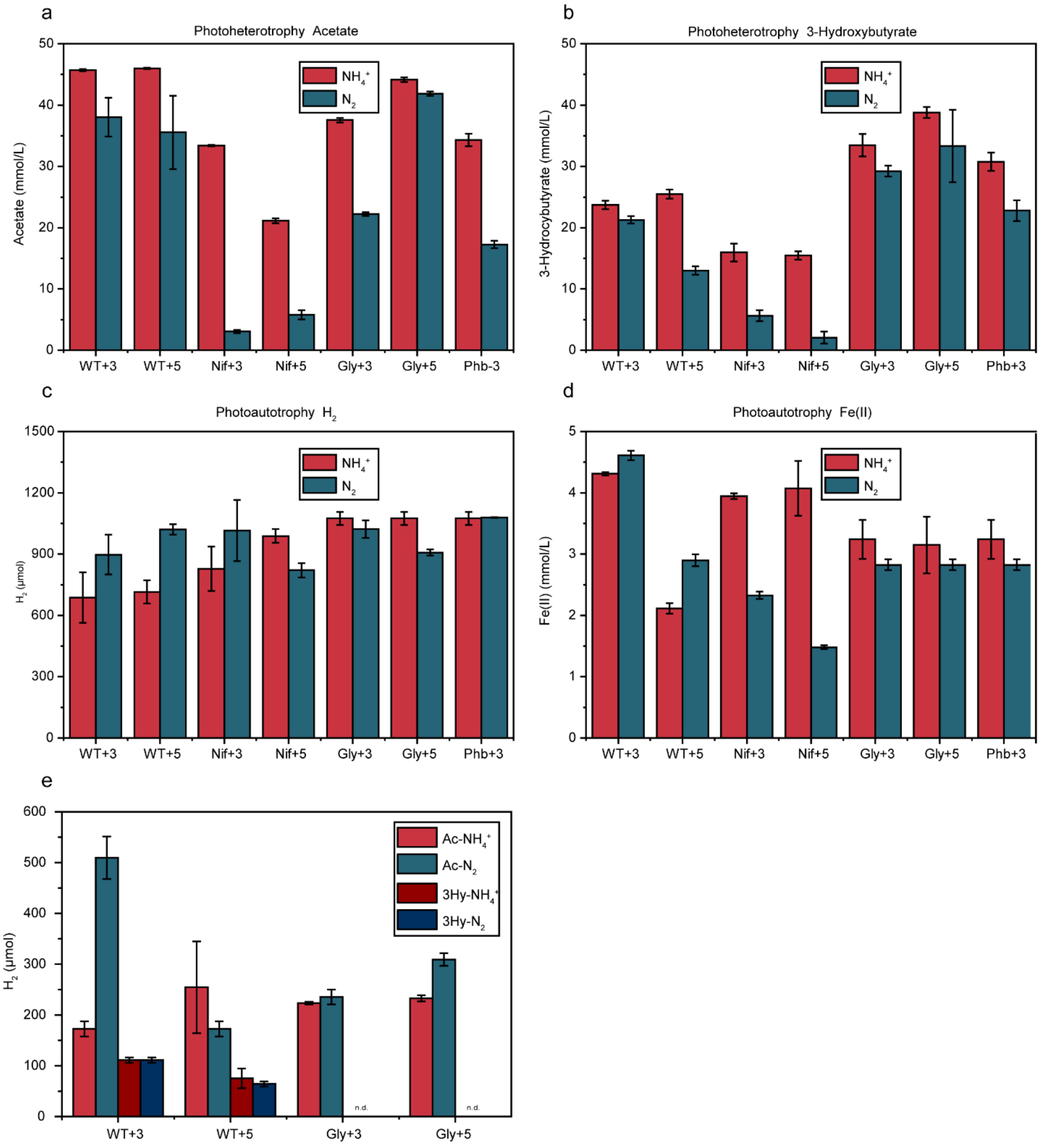


**Supplementary Figure 2 (a-e). Substrates consumption and hydrogen production under photoheterotrophic conditions.** (a-d) The electron donor consumption when TIE-1 was cultured with ammonium ($NH_4^+$, red) or dinitrogen gas ($N_2$, blue); (a) acetate (photoheterotrophy); (b) 3-hydroxybutyrate (photoheterotrophy); (c) hydrogen ($H_2$) (photoautotrophy); (d) ferrous iron (Fe(II)) (photoautotrophy); (e). $H_2$ production of WT+3/WT+5 and

Gly+3/Gly+5 mutant under photoheterotrophic conditions. Carbon dioxide was present in all conditions. $CO_2$: carbon dioxide; WT+3: wild type with 3-gene cassette; WT+5: wild type with 5-gene cassette; Nif+3: nitrogenase knockout t with 3-gene cassette; Nif+5: nitrogenase knockout with 5-gene cassette; Gly+3: glycogen synthase knockout with 3-gene cassette; Gly+5: glycogen synthase knockout with 5-gene cassette; Phb+3: hydroxybutyrate polymerase knockout with 3-gene cassette, n.d. (non-detectable).

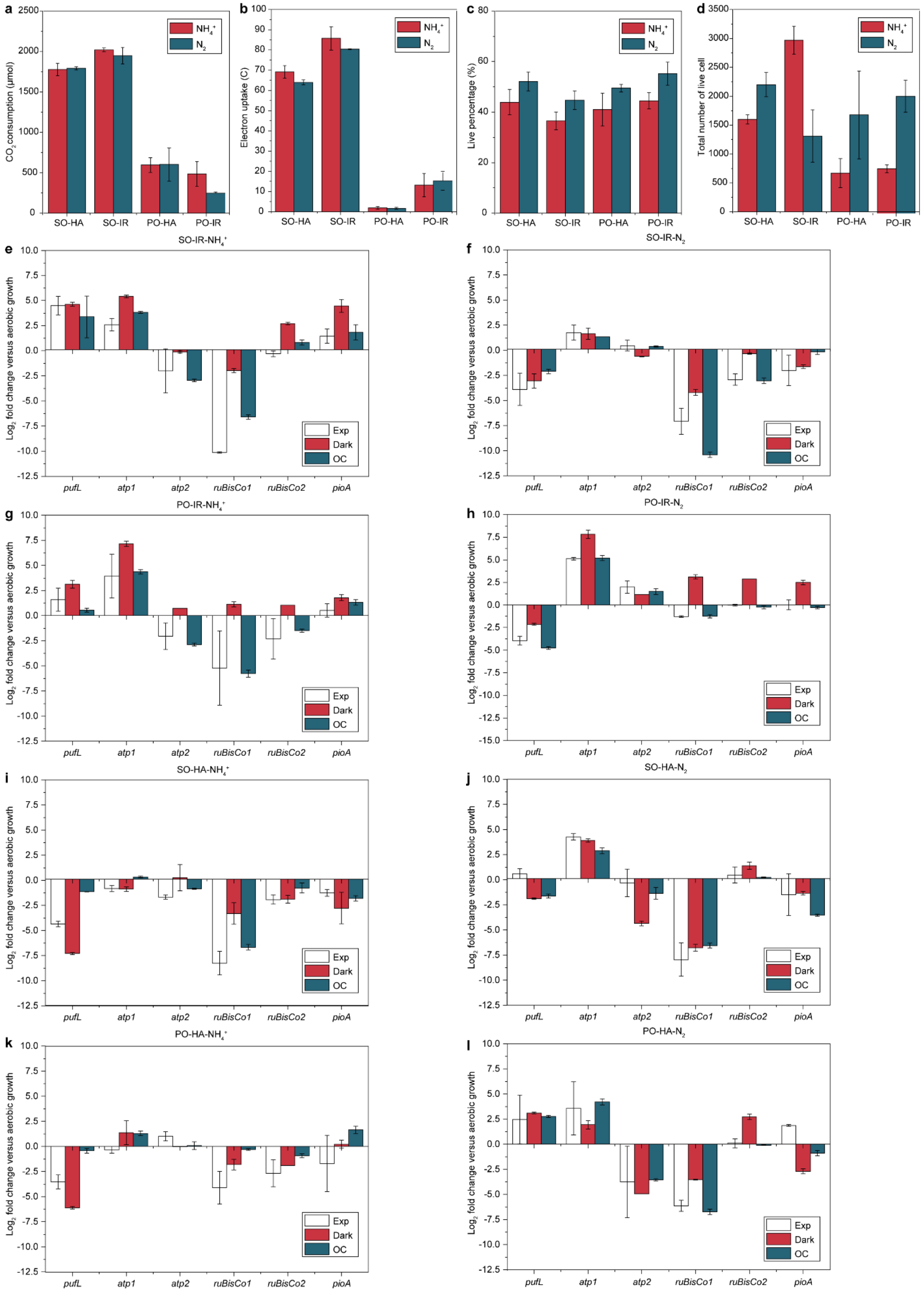
a
b
c
d
$CO_2$ consumption (μmol)
Electron uptake (C)
Live percentage (%)
Total number of live cell
$NH_4^+$
$N_2$
SO-HA
SO-IR
PO-HA
PO-IR
e
f
g
h
i
j
k
l
SO-IR-$NH_4^+$
SO-IR-$N_2$
PO-IR-$NH_4^+$
PO-IR-$N_2$
SO-HA-$NH_4^+$
SO-HA-$N_2$
PO-HA-$NH_4^+$
PO-HA-$N_2$
$Log_2$ fold change versus aerobic growth
Exp
Dark
OC
pufL
atp1
atp2
ruBisCo1
ruBisCo2
pioA

**Supplementary Figure 3. Carbon consumption, current uptake, and cell viability on the electrode and from each reactor setup.** **a.** Carbon dioxide consumption **b.** Current uptake **c.** The percentage of live cells **d.** The total number of live cells in the presence of ammonium ($NH_4^+$, red) or dinitrogen gas ($N_2$, blue) with various light and electricity sources. **e-l:** mRNA log2 fold change of photosynthetic reaction center (*pufL*), ATP synthase homologs (*atp*1, *atp*2), ribulose-1,5-bisphosphate carboxylase/oxygenase (*ruBisCo*1 and *roBisCo*2), and pio operon (*pioA)* in Nif mutant with 5-gene using different platforms**. e.** SO-IR-$NH_4^+$ **f.** SO-IR-$N_2$ **g.** PO-IR-$NH_4^+$ **h.** PO-IR-$N_2$ **i.** SO-HA-$NH_4^+$ **j.** SO-HA-$N_2$ **k.** PO-HA-$NH_4^+$ **l.** PO-HA-$N_2$ Data are means ± SD (standard deviation) of three biological replicates. SO: using electricity generated by a solar panel; HA: using halogen light as the light source. PO: using electricity from potentiostat as the electricity source; IR: using infrared light as the light source. $NH_4^+$: ammonium; $N_2$: dinitrogen gas. Exp: WT TIE-1 with illumination and closed circuit passing current; Dark: dark control group using WT TIE-1 without illumination; OC: open circuit control, WT TIE-1 with no electricity poised on the electrode.

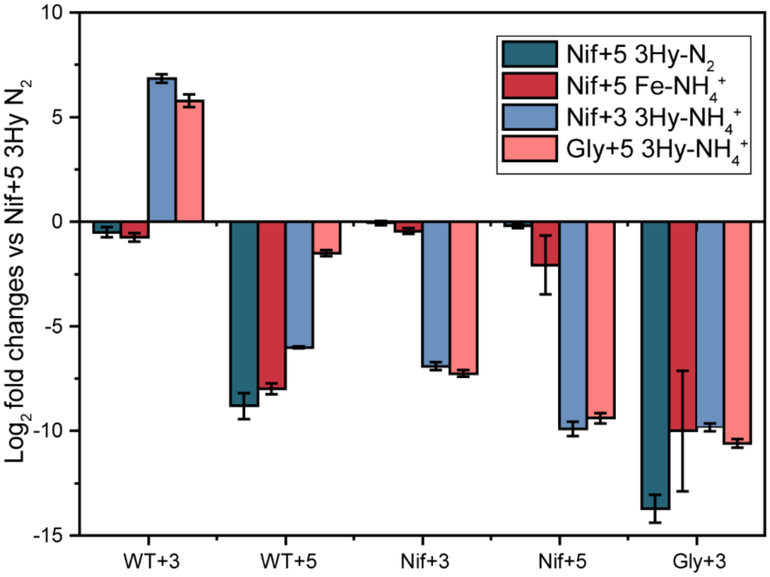


**Supplementary Figure 4. mRNA $log_2$ fold change of butanol synthesis genes (*phaJ*, *ter*, *adhE*2, *phaA, and phaB*).** Fold change was calculated using Nif mutant with 5-gene on 3Hy-$NH_4$ as the reference. The RT-qPCR data are means ± SD (standard deviations) of three biological replicates. Nif+3: nitrogenase knockout t with 3-gene cassette; Nif+5: nitrogenase knockout with 5-gene cassette; Gly+5: glycogen synthase knockout with 5-gene cassette; Phb+3: hydroxybutyrate polymerase knockout with 3-gene cassette, n.d. (non-detectable).

3Hy-$N_2$: using 3hydroxybutyrate as major carbon/electron source and dinitrogen gas ($N_2$) as the nitrogen source. Fe-$NH_4^+$: using carbon dioxide ($CO_2$) as carbon source, ferrous iron as electron source and ammonium ($NH_4^+$) as the nitrogen source. 3Hy-$NH_4^+$: using 3-hydroxybutyrate as major carbon/electron source and $NH_4^+$ as the nitrogen source. $CO_2$ was present in all conditions.

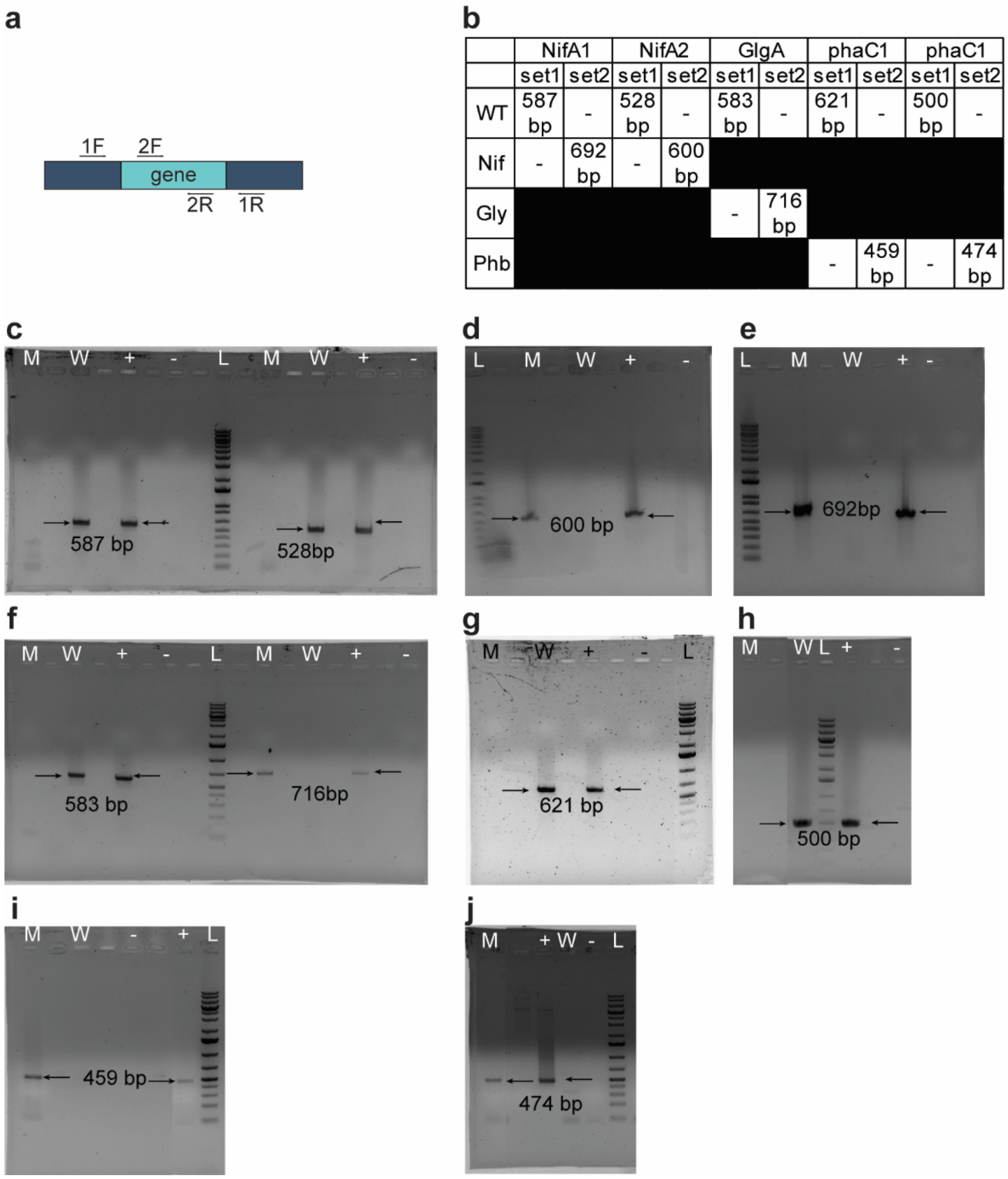


**Supplementary Figure 5. PCR check for mutants. a.** Schematic view of two different primer sets. **b.** Table of expected band sizes **c-e:** Nif mutant. **c.** To the left of ladder: primer set 2 for *nifA*1 (Rpal_1624). To the right of the ladder: primer set 2 for *nifA*2 (Rpal_5113). **d.** primer set 1 for *nifA*1 (Rpal_1624). **e.** Primer set 1 for *nifA*2 (Rpal_5113). **f.** Gly mutant. To the left of the ladder: primer set 2 for *glgA* (Rpal_0386). To the right of the ladder: primer set 1 for *glgA* (Rpal_0386). **g-j:** Phb mutant. **g.** Primer set 2 for *phaC*1 (Rpal_2780). **h.** primer set 2 for *phaC*2 (Rpal_4722). **i.** Primer set 1 for *phaC*1 (Rpal_2780). **j.** Primer set 1 for *phaC*2 (Rpal_4722). Genomic DNA from WT or TIE- 1 mutants was used as a PCR template. Depending on the primer set, either WT(W) or mutant (M) genomic DNA was used as the positive control (+). Autoclaved Mili Q water was used as negative control (-). In panel **c-f,** L: Thermal Fisher 1kb plus DNA ruler. In panel **g-j,** L**:** GeneRuler 1kb plus DNA ruler. M: mutant; W: Wild type; +: positive control; -: negative control.

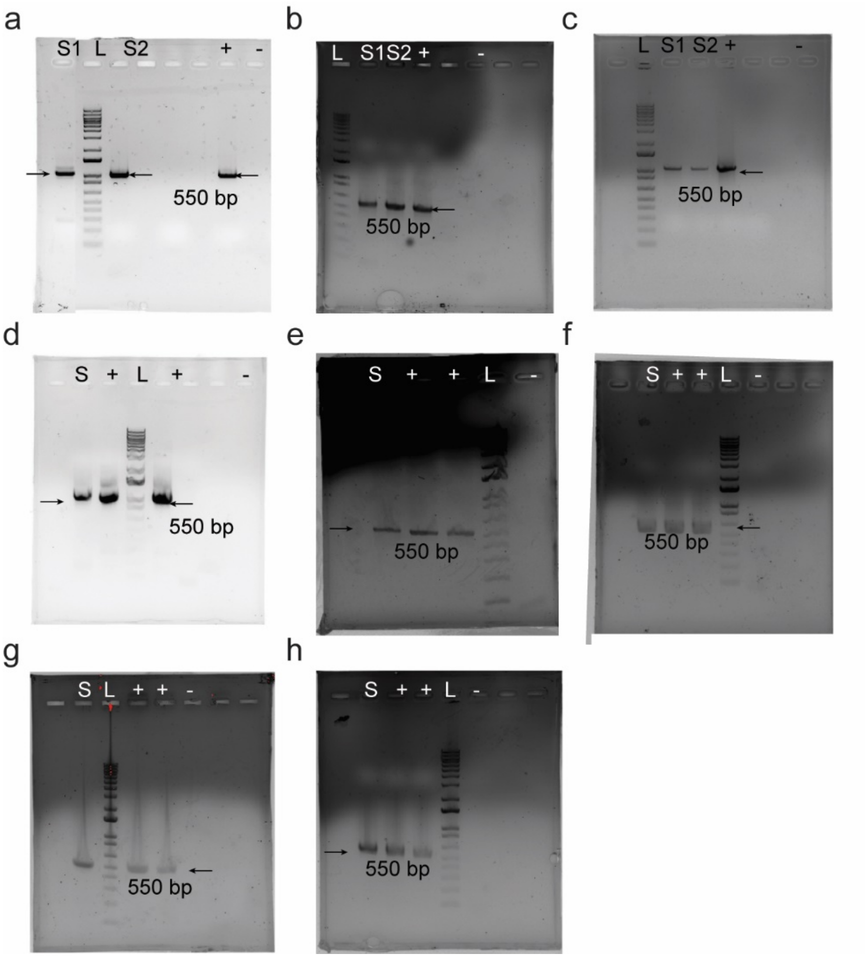


**Supplementary Figure 6. PCR checks of inoculum for *n*-butanol cassettes. A-D:** Inoculum for photoheterotrophic and photoautotrophic conditions using hydrogen or ferrous iron as electron donor **a.** WT. S1: WT with 3-gene cassette, S2: Wild Type with 5-gene cassette. **b.** Nif mutant. S1: Nif mutant with 3-gene cassette; S2Nif mutant with 5-gene cassette. **c.** Gly mutant. S1: Gly mutant with 3-gene cassette; S2: Gly mutant with 5-gene cassette. **d.** Phb mutant. S: Phb mutant with 3-gene cassette. **e-h:** Inoculums of Nif mutant with 5-gene cassette inoculum for photoelectroautotrophy. **e.** Photoelectroautotrophy using electricity from the solar panel under halogen light. Lane S: Nif mutant with 5-gene cassette. **f.** Photoelectroautotrophy using electricity from the solar panel under infrared light. S: Nif mutant with 5-gene cassette. **g.** Photoelectroautotrophy using electricity from potentiostat under halogen light. S: Nif mutant with 5-gene cassette. **h.** Photoelectroautotrophy using electricity from potentiostat under infrared light. S: Nif mutant with 5-gene cassette. A 550bp band is expected from both 3-gene and 5-gene cassette. Miniprep from inoculum was used as a PCR template. A plasmid with either the 3-gene cassette or the 5-gene cassette was used as a positive control. Autoclaved MiliQ water was used as negative control. L:1kb plus DNA ladder. S: sample; +: positive control; -: negative control.

**Supplementary Table 1 Strains, plasmids, incubation conditions and primers used in this study.**

**Supplementary Table 1-1 Incubation combinations used in this study (except photoelectroautotrophy)**

<table>
<tr><th colspan="2">Incubation condition</th><th>Nitrogen Sources</th><th>Electron Sources</th><th>Carbon Sources</th></tr>
<tr><td>1</td><td>Ac-$NH_4^+$</td><td>Ammonium ($NH_4^+$)</td><td rowspan="2">Acetate (Ac)</td><td rowspan="2">Acetate</td></tr>
<tr><td>2</td><td>Ac-$N_2$</td><td>Dinitrogen gas ($N_2$)</td></tr>
<tr><td>3</td><td>3Hy-$NH_4^+$</td><td>Ammonium</td><td rowspan="2">3-hydroxybutyrate (3Hy)</td><td rowspan="2">3-hydroxybutyrate</td></tr>
<tr><td>4</td><td>3Hy-$N_2$</td><td>Dinitrogen gas</td></tr>
<tr><td>5</td><td>$H_2$-$NH_4^+$</td><td>Ammonium</td><td rowspan="2">Hydrogen ($H_2$)</td><td rowspan="4">Carbon Dioxide</td></tr>
<tr><td>6</td><td>$H_2$-$N_2$</td><td>Dinitrogen gas</td></tr>
<tr><td>7</td><td>Fe(II)-$NH_4^+$</td><td>Ammonium</td><td rowspan="2">Ferrous Iron (Fe(II))</td></tr>
<tr><td>8</td><td>Fe(II)-$N_2$</td><td>Dinitrogen gas</td></tr>
</table>

**Supplementary Table 1-2 Incubation time of all cultures**

<table>
<tr><th></th><th>Ac-$NH_4^+$</th><th>Ac-$N_2$</th><th>3Hy-$NH_4^+$</th><th>3Hy-$N_2$</th><th>$H_2$-$NH_4^+$</th><th>$H_2$-$N_2$</th><th>Fe-$NH_4^+$</th><th>Fe-$N_2$</th><th>EU</th></tr>
<tr><td>WT+3</td><td colspan="4" rowspan="2">10 days</td><td colspan="4" rowspan="2">13 days</td><td rowspan="3">n.a.</td></tr>
<tr><td>WT+5</td></tr>
<tr><td>Nif+3</td><td rowspan="2">13days</td><td rowspan="2">16 days</td><td rowspan="2">13 days</td><td rowspan="2">16 days</td><td rowspan="2">16 days</td><td rowspan="2">19 days</td><td rowspan="2">16 days</td><td rowspan="2">19 days</td></tr>
<tr><td>Nif+5</td><td>10 days</td></tr>
<tr><td>Gly+3</td><td colspan="4" rowspan="3">10 days</td><td colspan="4" rowspan="3">13 days</td><td rowspan="3">n.a.</td></tr>
<tr><td>Gly+5</td></tr>
<tr><td>Phb+3</td></tr>
</table>

Ac: acetate; 3Hy: 3-hydroxybutyrate; $H_2$: hydrogen; Fe(II): ferrous iron; $CO_2$: carbon dioxide; $NH_4^+$ : ammonium; $N_2$: dinitrogen gas EU: photoelectroautotrophy WT+3: WT with 3-gene cassette; WT+5: WT with 5-gene cassette; Nif+3: *ΔnifA1ΔnifA2* with 3-gene cassette; Nif+5: *ΔnifA1ΔnifA2* with 5-gene cassette; Gly+3: *ΔglgA* with 3-gene cassette; Gly+5: *ΔglgA* with 5-gene cassette; Phb+3: *ΔphaC1ΔphaC2* with 3-gene cassette; n.d. :non-detectable; n. a. :non-applicable due to non-detectable results

**Supplementary Table 1-3 Final optical density ($OD_{660}$) of all the construct under *n*-butanol producing conditions.**

| | Ac-$NH_4^+$ | | Ac-$N_2$ | | 3Hy-$NH_4^+$ | | 3Hy-N2 | | $H_2$-$NH4^+$ | | $H_2$-$N_2$ | | Fe-$NH4^+$ | | Fe-$N_2$ | |
|---|---|---|---|---|---|---|---|---|---|---|---|---|---|---|---|---|
| | ave | std | ave | std | ave | std | ave | std | ave | std | ave | std | ave | std | ave | std |
| WT-3 | 4.13 | 0.85 | 3.98 | 0.25 | 2.71 | 0.32 | 2.62 | 0.52 | 1.23 | 0.09 | 0.99 | 0.16 | 1.22 | 0.11 | 1.14 | 0.28 |
| WT-5 | 5.14 | 0.49 | 1.52 | 0.32 | 3.61 | 0.19 | 1.44 | 0.08 | 1.67 | 0.16 | 1.53 | 0.02 | 1.89 | 0.05 | 1.73 | 0.10 |
| Nif-3 | 4.33 | 0.21 | 2.20 | 0.04 | 2.46 | 0.21 | 0.86 | 0.03 | 1.72 | 0.15 | 1.78 | 0.10 | 1.17 | 0.31 | 1.15 | 0.07 |
| Nif-5 | 3.30 | 0.44 | 1.12 | 0.02 | 2.92 | 0.17 | 1.05 | 0.03 | 1.85 | 0.05 | 1.20 | 0.19 | 1.73 | 0.06 | 0.74 | 0.05 |
| Gly-3 | 7.46 | 1.72 | 3.96 | 0.87 | 7.24 | 0.97 | 4.04 | 0.72 | 1.04 | 0.32 | 0.78 | 0.11 | 0.83 | 0.09 | 0.77 | 0.05 |
| Gly-5 | 4.86 | 0.60 | 2.47 | 0.31 | 3.26 | 0.18 | 1.86 | 0.08 | 1.41 | 0.07 | 1.54 | 0.27 | 1.56 | 0.11 | 1.44 | 0.08 |
| Phb-3 | 3.93 | 0.45 | 1.99 | 0.68 | 5.78 | 0.37 | 1.68 | 0.45 | 2.02 | 0.09 | 2.00 | 0.45 | 1.40 | 0.06 | 2.05 | 0.15 |

ave: average, std: standard deviation Ac: acetate; 3Hy: 3-hydroxybutyrate; $H_2$: hydrogen; Fe(II): ferrous iron; $CO_2$: carbon dioxide; $NH_4^+$: ammonium; $N_2$: dinitrogen gas EU: photoelectroautotrophy WT+3: WT with 3-gene cassette; WT+5: WT with 5-gene cassette; Nif+3: *ΔnifA1ΔnifA2* with 3-gene cassette; Nif+5: *ΔnifA1ΔnifA2* with 5-gene cassette; Gly+3: *ΔglgA* with 3-gene cassette; Gly+5: *ΔglgA* with 5-gene cassette; Phb+3: *ΔphaC1ΔphaC2* with 3-gene cassette;

**Supplementary Table 1-4 Incubation combinations used under photoelectroautotrophy**

<table>
<tr><th colspan="2">Platforms</th><th>Nitrogen Sources</th><th>Light Sources</th><th>Electricity Sources</th></tr>
<tr><td rowspan="2">1</td><td>SO-HA-$NH_4^+$</td><td>Ammonium ($NH_4^+$)</td><td rowspan="2">Halogen light (HA)</td><td rowspan="4">Solar panel (SO)</td></tr>
<tr><td>SO-HA-$N_2$</td><td>Dinitrogen gas ($N_2$)</td></tr>
<tr><td rowspan="2">2</td><td>SO-IR-$NH_4^+$</td><td>Ammonium</td><td rowspan="2">Infrared light (IR)</td></tr>
<tr><td>SO-IR-$N_2$</td><td>Dinitrogen gas</td></tr>
<tr><td rowspan="2">3</td><td>PO-HA-$NH_4^+$</td><td>Ammonium</td><td rowspan="2">Halogen light</td><td rowspan="4">Potentiostat (PO)</td></tr>
<tr><td>PO-HA-$N_2$</td><td>Dinitrogen gas</td></tr>
<tr><td rowspan="2">4</td><td>PO-IR-$NH_4^+$</td><td>Ammonium</td><td rowspan="2">Infrared light</td></tr>
<tr><td>PO-IR-$N_2$</td><td>Dinitrogen gas</td></tr>
</table>

**Supplementary Table 1-5 Strains used in this study.**

| Strains | Relevant genotypes of *R. palustris* TIE-1 | Plasmid | Reference |
|---|---|---|---|
| AB437 | Wild type (WT) | None | [47] |
| AB647 | Δ*phaC1*Δ*phaC2* (Phb) | None | This study |
| AB133 | Δ*nifA*1Δ*nifA*2 (Nif) | None | This study |
| AB145 | Δ*glgA* (Gly) | None | This study |
| AB147 | Wild type (WT) | pAB675 | This study |
| AB153 | Δ*phaC1*Δ*phaC2* (Phb) | pAB675 | This study |
| AB149 | Δ*nifA*1Δ*nifA*2 (Nif) | pAB675 | This study |
| AB151 | Δ*glgA* (Gly) | pAB675 | This study |
| AB148 | Wild type (WT) | pAB744 | This study |
| AB150 | Δ*nifA*1Δ*nifA*2 (Nif) | pAB744 | This study |
| AB152 | Δ*glgA* (Gly) | pAB744 | This study |

**Supplementary Table 1-6. Primers for constructing and sequencing the plasmids used for generating mutants**

| Primer name | Primer sequence |
|---|---|
| Rpal_51131kbDnFwXbaI | CATACTCTAGAAGCAGATCATCGTGGTGTCTTG |
| Rpal_51131kbDnBamHIRev | ATCAGGATCCCGCGGTCTCGGTGACCAGCTC |
| Rpal_51131kbUpFwSacI | TCATGAGCTCCAGAAGACGCTGGTGCTGAC |
| Rpal_51131kbupRevXbaI | GACTCTAGACATAGCTGGTCTCCATCGCTC |
| Rpal_1624 1Kb Up Spe I Fw NifA | TAGACTACTAGTCGTTTACAGCTCCGATCCGAATG |
| Rpal_1624 1Kb Up NifA BamHI Rev | ATACTAGGATCCGTCGGCTTCAGGACATGGTCG |
| Rpal_16241kbDnXbaIFw | CAGCTCTAGAATCATCAGACAAGGCGCGAC |
| Rpal_16241kbDnBamHlIRev | TCATGGATCCATTGGCAAGCGCATCACCCGGACC |
| Rpal_2780 1Kb Up Fw | CATATGACTAGTGAGTGTCCTTCAGCTTCTCCAGGA |
| Rpal_2780 1Kb Up Rev | CATATGGGATCCGAATCAACACTACAGTCCGGT |
| Rpal_2780 1Kb Dn Fw | CATATGGGATCCTGAACGACGCGCGGCGGCGAAGC |
| Rpal_2780 1Kb Dn Rev | CATATGCCCGGGACGGTGAGCACCGAATTCGCCTG |
| Rpal_4722 1Kb Up Fw | CATATGGCGGCCGCCGCGTGTCGTCTCAGCATTGCG |
| Rpal_4722 1Kb Up Rev | CATATGGGATCCCATCACCTCGTCGCGGCCGTC |
| Rpal_4722 1Kb Dn Fw | CATATGGGATCCTGAGCGGTCTGCGGCAACGCCGC |
| Rpal_4722 1Kb Dn Rev | CATATGCTGCAGTGGCCGACGACACCAACGAGCT |
| Gly (Rpal_0386) UP XbaI F | GCTATATCTAGAAGCGCAACGAGAGCTTCGACATTCTGCC |
| Gly (Rpal0386) UP BamHI R | ATATATGGATCCGCGTCGAGCTTCTCGATCATCGC |
| Gly (Rpal_0386) DN BamHI F | ATATATGGATCCCAAGCTGCGCACATCATCCCA |
| Gly (Rpal_0386) DN XhoI R | GCATATCTCGAGTCTGATCATGGAGCCGGCTTG |
| Seq gly (Rpal_0386) F | GAGAACACTGTGGTGCT |

**Supplementary Table 1-7. Primers used for checking the mutants**

| Primer name | Primer sequence |
|---|---|
| PCR check Rpal_5113 set 1 F | GAGTGCTGACCTGAGCGAATAG |
| PCR check Rpal_5113 set 1R | CACTTGGTCGCGTCGATCACATAG |
| PCR check Rpal_5113 set 2 F | CGTTCGCACTTCCGGATGGAC |
| PCR check Rpal_5113 set 2 R | CTTGATGGTCTGGTTGCCGCCGAC |
| PCR check Rpal_1624 set 1 F | GAACCAGCTCGCGATCCATCTCAG |
| PCR check Rpal_1624 set 1R | GCCACGATGTAGACTTCCTGTGCCTTG |
| PCR check Rpal_1624 set 2 F | GAGCACCTTCTGGGCGAGCACGATC |
| PCR check Rpal_1624 set 2 R | ATGCCGATGGCCCAAAATTCCCG |
| PCR check Rpal_2780 set 1 F | CTCGCAACAATCGTCGCACTC |
| PCR check Rpal_2780 set 1R | AAGGATTGGCCTATACCG |
| PCR check Rpal_2780 set 2 F | CTTCCAGAACGAAATCATGCAGCTC |
| PCR check Rpal_2780 set 2 R | GGTGGCGTCGGAATTCCAGT |
| PCR check Rpal_4722 set 1 F | ATCCGCGGCTTGAGCAAGG |
| PCR check Rpal_4722 set 1R | TTGGCATCCCATTCCACG |
| PCR check Rpal_4722 set 2 F | GTCAAACTTCGCCCTCACCAATC |
| PCR check Rpal_4722 set 2 R | CGAGGCAATAGCCGACCG |
| PCR check Rpal_0386 set 1 F | TCACTTCGACAAGTCGTGCC |
| PCR check Rpal_0386 set 1 R | TAATCGTGACGATGGTCACC |
| PCR check Rpal_0386 set 2 F | AGGTCCACAGCTTCAACGAG |
| PCR check Rpal_0386 set 2 R | GTGTCGATGCCGTTGAGGAT |

**Supplementary Table 1-8 Plasmids used in this study.**

| Plasmid | Construct | Reference |
|---|---|---|
| pAB423 | Empty Vector | [48] |
| pAB675 | *phaJ, ter, adhE2* in pAB423 | This study |
| pAB744 | *phaJ, ter, adhE2, phaA, phaB* in pAB423 | This study |

**Supplementary Table 1-9. Primers for constructing the 3-gene and the 5-gene cassette**

| Primer name | Primer sequence |
|---|---|
| 1F | ATCGAATTCCGCTAGCTTCACGCTGCCGCA |
| 12F | GACCCGGCGTTTGCGGCGACCACGGCGTTC |
| 23F | GTCGCGCATCACCGCCGCCTTCGGCTACGG |
| 34F | ACTCGTATATCGGCCCGGAAGCGACCCAGG |
| 45F | GACCGCATCTAAATGCATGCAGGATGAGGA |
| 56F | GGACGCCGCCGTGAAGGCCGGCGCGCCGAA |
| 67F | TGCAGTCCGTCGAGAAGTCGGAGCTGTTCA |
| 78F | CTGTTCAAGCTGGGCTACGTCAACAAGATC |
| 89F | CCACGCGATCGAGGCGTATGTGTCGGTGAT |
| 12R | GAACGCCGTGGTCGCCGCAAACGCCGGGTC |
| 23R | CCGTAGCCGAAGGCGGCGGTGATGCGCGAC |
| 34R | CCTGGGTCGCTTCCGGGCCGATATACGAGT |
| 45R | TCCTCATCCTGCATGCATTTAGATGCGGTC |
| 56R | TTCGGCGCGCCGGCCTTCACGGCGGCGTCC |
| 67R | TGAACAGCTCCGACTTCTCGACGGACTGCA |
| 78R | GATCTTGTTGACGTAGCCCAGCTTGAACAG |
| 89R | ATCACCGACACATACGCCTCGATCGCGTGG |
| 10R | CGATCGATCGATCGATCTGCAGCTCCAAAA |
| 910F | TCTACAACACCCTCGACAAGATGAGCGAGC |
| 910R | CGATCGATCGATCGATCTGCAGCTCCAAAA |
| But XbaI NdeI F | CCGAGTCTAGACGTTTCATATGTCCGC |
| But EcoRI R | GCGCGAATTCTTAGATGCGGTCGAAGC |
| But EcoRI F | GCATGAATTCAGGATGAGGATCGTTTCGCATGAAGGT |

| | |
|---|---|
| But KpnI R | GTTAGGTACCGATCGATCGATCCATCTGCAGCTCC |
| phaA HindIII F | ATCGGCAAGCTTCTAACCAGGAGATGTCCATGTCGGA |
| phaB PstI R | ATATCTGCAGTCAAACCATGTATTGGCCGCCGTTGATGGTGA |

**Supplementary Table 1-10. Primers for sequencing the 3-gene and the 5-gene cassette**

| Primer name | Primer sequence |
|---|---|
| But seq 2 | GCGAGGACAAGCCAATCGCCACCCTCACCACCCGCATC |
| But seq 3 | GCCTACTCGTATATCGGCCCGGAAGCGACCCAGGCCCTC |
| But seq new 2 | AGTCACGGCCGAGGTGGAAG |
| But Seq new 3 | GCGTCCTGAAGCCGTTCG |
| But Seq 4 | CGGCATCATCGACCACGACGACAGCCTCGGCATCACCAAG |
| But Seq 5 | CGGGCCATACCTCGTCGCTGTATATCGACAGCCAGAAC |
| phaA phaB Seq2 | TGGTGCTGATGACCGCCAA |
| phaA phaB seq3 | TGGGACGTGAGTTCGTTCGA |

**Supplementary Table 1-11. Primers for q-RTPCR**

| Primer name | Primer sequence |
|---|---|
| qPCR phaJ F | CCTTTAAGCTGCCGGTGTTC |
| qPCR phaJ R | GTGGTGAGGGTGGCGATT |
| qPCR ter F | GCGCAACAATATCTGCCTGA |
| qPCR ter R | TGATGCGCTTCTTGGTGTAC |
| qPCR adhe2 F | ACCTGCTGTACGAGTATCCG |
| qPCR adhe2 R | CAGCTTCGGGAAATTGCAGA |
| qPCR phaA F | GGATGCCTTCAACGGTTACC |
| qPCR phaA R | CGCGAATTCATCCTGCTGAT |
| qPCR phaB F | TTTGAATTTCTCCGCTGCCG |
| qPCR phaB R | GGTATCGGTGCAGCAATCAG |
| TIE-1recAqRT-PCRFor | ATCGGCCAGATCAAGGAAC |
| TIE-1recAqRT-PCRRev | GAATTCGACCTGCTTGAACG |
| TIE-1clpXqRT-PCRFor | GGAGATCTGCAAGGTTCTCG |
| TIE-1clpXqRT-PCRRev | CCGCTTGTAGTGATTGTGGA |
| Km qPCR_F | CTCGTCCTGCAGTTCATTCA |
| Km qPCR_R | AGACAATCGGCTGCTCTGAT |
| Rpal_1747_rubisco1_qPCR_for | ACCAAGGACGACGAGAACAT |
| Rpal_1747_rubisco1_qPCR_rev | CATGCAGTATTGGAAGCGCT |
| Rpal_5122_rubisco2_qPCR_for | GGCGTATCTCAAGCTGTTCG |
| Rpal_5122_rubisco2_qPCR_rev | CGATGAAGCCACCGTTGATC |
| Rpal_1716_pufL_qPCR_for | GAGAAGAAATACCGCGTTCG |
| Rpal_1716_pufL_qPCR_rev | CCGAAGATCCCAACGTAGAA |
| Rpal_1057_atp1_qPCR_for | ATTCTGAACGCCATCGAAAC |
| Rpal_1057_atp1_qPCR_rev | GACGGTCGATTCACCAAGAT |

| | |
|---|---|
| Rpal_0171_atp2_qPCR_for | GACGATCGCGGAGTATTTTC |
| Rpal_0171_atp2_qPCR_rev | CAGGCTGGATAACTCGCTTC |
| Rpal_0817_pioA_qPCR_for | AAATTTCGACGACACCATCGA |
| Rpal_0817_pioA_qPCRrev | CTTGGCGGCGAGGATCT |

116

**Supplementary Table 2. *p*-value**.

For butanol and acetone, all means and standard deviations are in mg/L, for $H_2$ or $CO_2$ generations/consumptions all means and standard deviations are in μmol for conversions all the means and standard deviations are in percentage. *P*<0.05 is considered significant.

**Supplementary Table 2-1.** *p*-value between *n*-butanol production from different carbon and electron sources when incubated with $NH_4^+$

| | WT+3 | | | WT+5 | | | Nif+3 | | | Nif+5 | | | Gly+3 | | | Gly+5 | | | Phb+3 | | |
|---|---|---|---|---|---|---|---|---|---|---|---|---|---|---|---|---|---|---|---|---|---|
| Substrates | Mean | s.e.m. | P-value | Mean | s.e.m. | P-value | Mean | s.e.m. | P-value | Mean | s.e.m. | P-value | Mean | s.e.m. | P-value | Mean | s.e.m. | P-value | Mean | s.e.m. | P-value |
| Ac + $CO_2$ | 2.26 | 0.05 | n.a. | 1.29 | 0.28 | n.a. | 2.76 | 0.23 | n.a. | 3.73 | 0.95 | n.a. | 0.13 | 0.02 | n.a. | n.d. | n.a. | n.a. | 0.67 | 0.01 | n.a. |
| 3Hy + $CO_2$ | 1.40 | 0.22 | 0.02 | 2.17 | 0.05 | 0.04 | 3.60 | 0.15 | 0.04 | 4.98 | 0.87 | 0.39 | 1.22 | 0.03 | 0.00 | n.d. | n.a. | n.a. | 0.68 | 0.07 | 0.97 |
| $H_2$ + $CO_2$ | 1.46 | 0.38 | 0.10 | 1.95 | 0.47 | 0.30 | 2.36 | 0.34 | 0.39 | 3.36 | 0.10 | 0.72 | 0.28 | 0.03 | 0.02 | n.d. | n.a. | n.a. | 0.20 | 0.07 | 0.02 |
| Fe(II) + $CO_2$ | 0.30 | 0.05 | 0.00 | 0.28 | 0.03 | 0.02 | 0.71 | 0.04 | 0.00 | 0.55 | 0.03 | 0.03 | n.d. | n.a. | n.a. | n.d. | n.a. | n.a. | n.d. | n.a. | n.a. |
| 3Hy + $CO_2$ | 1.40 | 0.22 | n.a. | 2.17 | 0.05 | n.a. | 3.60 | 0.15 | n.a. | 4.98 | 0.87 | n.a. | 1.22 | 0.03 | n.a. | n.d. | n.a. | n.a. | 0.68 | 0.07 | n.a. |
| $H_2$ + $CO_2$ | 1.46 | 0.38 | 0.90 | 1.95 | 0.47 | 0.67 | 2.36 | 0.34 | 0.03 | 3.36 | 0.10 | 0.14 | 0.28 | 0.03 | 0.00 | n.d. | n.a. | n.a. | 0.20 | 0.07 | 0.01 |
| Fe(II) + $CO_2$ | 0.30 | 0.05 | 0.01 | 0.28 | 0.03 | 0.00 | 0.71 | 0.04 | 0.00 | 0.55 | 0.03 | 0.01 | n.d. | n.a. | n.a. | n.d. | n.a. | n.a. | n.d. | n.a. | n.a. |
| $H_2$ + $CO_2$ | 1.46 | 0.38 | n.a. | 1.95 | 0.47 | n.a. | 2.36 | 0.34 | n.a. | 3.36 | 0.10 | n.a. | 0.28 | 0.03 | n.a. | n.d. | n.a. | n.a. | 0.20 | 0.07 | n.a. |
| Fe(II) + $CO_2$ | 0.30 | 0.05 | 0.04 | 0.28 | 0.03 | 0.02 | 0.71 | 0.04 | 0.01 | 0.55 | 0.03 | 0.00 | n.d. | n.a. | n.a. | n.d. | n.a. | n.a. | n.d. | n.a. | n.a. |

For row 1~4, the *p*-value is between the substrate combination in the row 1 and the respective row. For row 5~7, the *p*-value is between the substrate combination in the row 5 and the respective row. For row 8~9, the *p*-value is between the substrate combination in the row 8 and the respective row.

**Supplementary Table 2-2.** *p*-value between *n*-butanol production from different carbon and electron sources when incubated with $N_2$

| | WT+3 | | | WT+5 | | | Nif+3 | | | Nif+5 | | | Gly+3 | | | Gly+5 | | | Phb+3 | | |
|---|---|---|---|---|---|---|---|---|---|---|---|---|---|---|---|---|---|---|---|---|---|
| Substrates | Mean | s.e.m. | P-value | Mean | s.e.m. | P-value | Mean | s.e.m. | P-value | Mean | s.e.m. | P-value | Mean | s.e.m. | P-value | Mean | s.e.m. | P-value | Mean | s.e.m. | P-value |
| Ac + $CO_2$ | 1.85 | 0.32 | n.a. | 1.18 | 0.20 | n.a. | 2.17 | 0.24 | n.a. | 2.70 | 0.27 | n.a. | 0.19 | 0.05 | n.a. | n.d. | n.a. | n.a. | 0.28 | 0.03 | n.a. |
| 3Hy + $CO_2$ | 2.26 | 0.35 | 0.44 | 2.64 | 0.11 | 0.00 | 2.78 | 0.30 | 0.19 | 2.02 | 0.42 | 0.24 | 0.88 | 0.13 | 0.01 | 0.67 | 0.06 | n.a. | 0.37 | 0.06 | 0.27 |
| $H_2$ + $CO_2$ | 2.31 | 0.05 | 0.23 | 2.03 | 0.09 | 0.02 | 3.09 | 0.25 | 0.06 | 2.43 | 0.27 | 0.55 | 0.30 | 0.01 | 0.10 | 0.24 | 0.02 | n.a. | 0.20 | 0.03 | 0.13 |
| Fe(II) + $CO_2$ | 0.30 | 0.04 | 0.01 | 0.22 | 0.03 | 0.01 | 0.79 | 0.03 | 0.00 | 0.57 | 0.07 | 0.00 | n.d. | n.a. | n.a. | n.d. | n.a. | n.a. | n.d. | n.a. | n.a. |
| 3Hy + $CO_2$ | 2.26 | 0.35 | n.a. | 2.64 | 0.11 | n.a. | 2.78 | 0.30 | n.a. | 2.02 | 0.42 | n.a. | 0.88 | 0.13 | n.a. | 0.67 | 0.06 | n.a. | 0.37 | 0.06 | n.a. |
| $H_2$ + $CO_2$ | 2.31 | 0.05 | 0.89 | 2.03 | 0.09 | 0.01 | 3.09 | 0.25 | 0.47 | 2.43 | 0.27 | 0.53 | 0.30 | 0.01 | 0.01 | 0.24 | 0.02 | 0.00 | 0.20 | 0.03 | 0.07 |
| Fe(II) + $CO_2$ | 0.30 | 0.04 | 0.01 | 0.22 | 0.03 | 0.00 | 0.79 | 0.03 | 0.00 | 0.57 | 0.07 | 0.03 | n.d. | n.a. | n.a. | n.d. | n.a. | n.a. | n.d. | n.a. | n.a. |
| $H_2$ + $CO_2$ | 2.31 | 0.05 | n.a. | 2.03 | 0.09 | n.a. | 3.09 | 0.25 | n.a. | 2.43 | 0.27 | n.a. | 0.30 | 0.01 | n.a. | 0.24 | 0.02 | n.a. | 0.20 | 0.03 | n.a. |
| Fe(II) + $CO_2$ | 0.30 | 0.04 | 0.00 | 0.22 | 0.03 | 0.00 | 0.79 | 0.03 | 0.00 | 0.57 | 0.07 | 0.00 | n.d. | n.a. | n.a. | n.d. | n.a. | n.a. | n.d. | n.a. | n.a. |

For row 1~4, the *p*-value is between the substrate combination in the row 1 and the respective row. For row 5~7, the *p*-value is between the substrate combination in the row 5 and the respective row. For row 8~9, the *p*-value is between the substrate combination in the row 8 and the respective row.

**Supplementary Table 2-3.** *p*-value between *n*-butanol production from WT and mutants

| | | Ac-$NH_4^+$ | | | Ac-$N_2$ | | | 3Hy-$NH_4^+$ | | | 3Hy-$N_2$ | | | $H_2$-$NH_4^+$ | | | $H_2$-$N_2$ | | | Fe(II)-$NH_4^+$ | | | Fe(II)-$N_2$ | | |
|---|---|---|---|---|---|---|---|---|---|---|---|---|---|---|---|---|---|---|---|---|---|---|---|---|---|
| | Genotype | Mean | s.e.m. | P-value | Mean | s.e.m. | P-value | Mean | s.e.m. | P-value | Mean | s.e.m. | P-value | Mean | s.e.m. | P-value | Mean | s.e.m. | P-value | Mean | s.e.m. | P-value | Mean | s.e.m. | P-value |
| **3-gene** | WT | 2.26 | 0.05 | n.a. | 1.85 | 0.32 | n.a. | 1.40 | 0.22 | n.a. | 2.26 | 0.35 | n.a. | 1.46 | 0.38 | n.a. | 2.31 | 0.05 | n.a. | 0.30 | 0.05 | n.a. | 0.30 | 0.04 | n.a. |
| | Nif | 2.76 | 0.23 | 0.10 | 2.17 | 0.24 | 0.47 | 3.60 | 0.15 | 0.00 | 2.78 | 0.30 | 0.32 | 2.36 | 0.34 | 0.15 | 3.09 | 0.25 | 0.04 | 0.71 | 0.04 | 0.00 | 0.79 | 0.03 | 0.00 |
| | Gly | 0.13 | 0.02 | 0.00 | 0.19 | 0.05 | 0.01 | 1.22 | 0.03 | 0.56 | 0.88 | 0.13 | 0.02 | 0.28 | 0.03 | 0.04 | 0.30 | 0.01 | 0.00 | n.d. | n.a. | n.a. | n.d. | n.a. | n.a. |
| | Phb | 0.67 | 0.01 | 0.00 | 0.28 | 0.03 | 0.01 | 0.68 | 0.07 | 0.03 | 0.37 | 0.06 | 0.01 | 0.20 | 0.07 | 0.03 | 0.20 | 0.03 | 0.00 | n.d. | n.a. | n.a. | n.d. | n.a. | n.a. |
| **5-gene** | WT | 1.29 | 0.28 | n.a. | 1.18 | 0.20 | n.a. | 2.17 | 0.05 | n.a. | 2.64 | 0.11 | n.a. | 1.95 | 0.47 | n.a. | 2.03 | 0.09 | n.a. | 0.28 | 0.03 | n.a. | 0.22 | 0.03 | n.a. |
| | Nif | 3.73 | 0.95 | 0.07 | 2.70 | 0.27 | 0.01 | 4.98 | 0.87 | 0.03 | 2.02 | 0.42 | 0.22 | 3.36 | 0.10 | 0.04 | 2.43 | 0.27 | 0.19 | 0.55 | 0.03 | 0.00 | 0.57 | 0.07 | 0.01 |
| | Gly | n.d. | n.a. | n.a. | n.d. | n.a. | n.a. | n.d. | n.a. | n.a. | 0.67 | 0.06 | 0.00 | n.d. | n.a. | n.a. | 0.24 | 0.02 | 0.00 | n.d. | n.a. | n.a. | n.d. | n.a. | n.a. |

For row 1~4, the *p*-value is between the substrate combination in the row 1 and the respective row. For row 5~7, the *p*-value is between the substrate combination in the row 5 and the respective row.

**Supplementary Table 2-4.** *p*-value between acetone production from WT and mutants

| | | Ac-$NH_4^+$ | | | Ac-$N_2$ | | | 3Hy-$NH_4^+$ | | | 3Hy-$N_2$ | | | $H_2$-$NH_4^+$ | | | $H_2$-$N_2$ | | | Fe(II)-$NH_4^+$ | | | Fe(II)-$N_2$ | | |
|---|---|---|---|---|---|---|---|---|---|---|---|---|---|---|---|---|---|---|---|---|---|---|---|---|---|
| | Genotype | Mean | s.e.m. | P-value | Mean | s.e.m. | P-value | Mean | s.e.m. | P-value | Mean | s.e.m. | P-value | Mean | s.e.m. | P-value | Mean | s.e.m. | P-value | Mean | s.e.m. | P-value | Mean | s.e.m. | P-value |
| 3-gene | WT | n.d. | n.a. | n.a. | n.d. | n.a. | n.a. | 80.09 | 3.13 | n.a. | 77.18 | 5.08 | n.a. | n.d. | n.a. | n.a. | n.d. | n.a. | n.a. | 0.78 | 0.11 | n.a. | 0.67 | 0.08 | n.a. |
| | Nif | 8.66 | 2.74 | n.a. | n.d. | n.a. | n.a. | 76.44 | 1.12 | 0.33 | 22.02 | 2.55 | 0.00 | n.d. | n.a. | n.a. | n.d. | n.a. | n.a. | 1.20 | 0.12 | 0.06 | 1.59 | 0.20 | 0.01 |
| | Gly | 19.35 | 1.73 | n.a. | 37.29 | 3.40 | n.a. | 99.80 | 4.67 | 0.02 | 192.84 | 4.82 | 0.00 | n.d. | n.a. | n.a. | 1.30 | 0.26 | n.a. | 2.13 | 0.65 | 0.11 | 3.01 | 0.65 | 0.02 |
| | Phb | 27.69 | 2.44 | n.a. | 25.60 | 2.54 | n.a. | 290.10 | 47.51 | 0.01 | 131.94 | 29.02 | 0.14 | 1.07 | 0.36 | n.a. | 1.55 | 0.63 | n.a. | n.d. | n.a. | n.a. | n.d. | n.a. | n.a. |
| 5-gene | WT | n.d. | n.a. | n.a. | 1.06 | 0.20 | n.a. | 107.39 | 3.74 | n.a. | 102.13 | 1.80 | n.a. | n.d. | n.a. | n.a. | n.d. | n.a. | n.a. | 0.79 | 0.16 | n.a. | 0.67 | 0.11 | n.a. |
| | Nif | n.d. | n.a. | n.a. | n.d. | n.a. | n.a. | 69.48 | 10.05 | 0.02 | 18.15 | 1.41 | 0.00 | n.d. | n.a. | n.a. | n.d. | n.a. | n.a. | n.d. | n.a. | n.a. | n.d. | n.a. | n.a. |
| | Gly | 8.50 | 1.93 | n.a. | 5.00 | 0.87 | 0.01 | 74.28 | 1.58 | 0.00 | 104.80 | 26.56 | 0.92 | 1.89 | 0.18 | n.a. | 1.47 | 0.08 | n.a. | 3.92 | 0.44 | 0.00 | 3.48 | 0.40 | 0.00 |

For row 1~4, the *p*-value is between the substrate combination in the row 1 and the respective row. For row 5~7, the *p*-value is between the substrate combination in the row 5 and the respective row.

**Supplementary Table 2-5.** *p*-value between acetone productions from different carbon and electron sources when incubated with $NH_4^+$

| | | WT+3 | | | WT+5 | | | Nif+3 | | | Nif+5 | | | Gly+3 | | | Gly+5 | | | Phb+3 | | |
|---|---|---|---|---|---|---|---|---|---|---|---|---|---|---|---|---|---|---|---|---|---|---|
| | Substrates | Mean | s.e.m. | P-value | Mean | s.e.m. | P-value | Mean | s.e.m. | P-value | Mean | s.e.m. | P-value | Mean | s.e.m. | P-value | Mean | s.e.m. | P-value | Mean | s.e.m. | P-value |
| Ac + $CO_2$ | Ac + $CO_2$ | n.d. | n.a. | n.a. | n.d. | n.a. | n.a. | 8.66 | 2.74 | n.a. | n.d. | n.a. | n.a. | 19.35 | 1.73 | n.a. | 8.50 | 1.93 | n.a. | 27.69 | 2.44 | n.a. |
| | 3Hy + $CO_2$ | 80.09 | 3.13 | n.a. | 107.39 | 3.74 | n.a. | 76.44 | 1.12 | 0.00 | 69.48 | 10.05 | n.a. | 99.80 | 4.67 | 0.00 | 74.28 | 1.58 | 0.00 | 290.10 | 47.51 | 0.01 |
| | $H_2$ + $CO_2$ | n.d. | n.a. | n.a. | n.d. | n.a. | n.a. | n.d. | n.a. | n.a. | n.d. | n.a. | n.a. | n.d. | n.a. | n.a. | 1.89 | 0.18 | 0.03 | 1.07 | 0.36 | 0.00 |
| | Fe(II) + $CO_2$ | 0.78 | 0.11 | n.a. | 0.79 | 0.16 | n.a. | 1.20 | 0.12 | 0.05 | n.d. | n.a. | n.a. | 2.13 | 0.65 | 0.00 | 3.92 | 0.44 | 0.08 | n.d. | n.a. | n.a. |
| 3Hy + $CO_2$ | 3Hy + $CO_2$ | 80.09 | 3.13 | n.a. | 107.39 | 3.74 | n.a. | 76.44 | 1.12 | n.a. | 69.48 | 10.05 | n.a. | 99.80 | 4.67 | n.a. | 74.28 | 1.58 | n.a. | 290.10 | 47.51 | n.a. |
| | $H_2$ + $CO_2$ | n.a. | n.a. | n.a. | n.a. | n.a. | n.a. | n.a. | n.a. | n.a. | n.a. | n.a. | n.a. | n.a. | n.a. | n.a. | 1.89 | 0.18 | 0.00 | 1.07 | 0.36 | 0.00 |
| | Fe(II) + $CO_2$ | 0.78 | 0.11 | 0.00 | 0.79 | 0.16 | 0.00 | 1.20 | 0.12 | 0.00 | n.a. | n.a. | n.a. | 2.13 | 0.65 | 0.00 | 3.92 | 0.44 | 0.00 | n.a. | n.a. | n.a. |
| $H_2$ + $CO_2$ | $H_2$ + $CO_2$ | n.a. | n.a. | n.a. | n.a. | n.a. | n.a. | n.a. | n.a. | n.a. | n.a. | n.a. | n.a. | n.a. | n.a. | n.a. | 1.89 | 0.18 | n.a. | 1.07 | 0.36 | n.a. |
| | Fe(II) + $CO_2$ | 0.78 | 0.11 | n.a. | 0.79 | 0.16 | n.a. | 1.20 | 0.12 | n.a. | n.a. | n.a. | n.a. | 2.13 | 0.65 | n.a. | 3.92 | 0.44 | 0.01 | n.a. | n.a. | n.a. |

For row 1~4, the *p*-value is between the substrate combination in the row 1 and the respective row. For row 5~7, the *p*-value is between the substrate combination in the row 5 and the respective row. For row 8~9, the *p*-value is between the substrate combination in the row 8 and the respective row.

**Supplementary Table 2-6.** *p*-value between acetone productions from different carbon and electron sources when incubated with $N_2$

| | | WT+3 | | | WT+5 | | | Nif+3 | | | Nif+5 | | | Gly+3 | | | Gly+5 | | | Phb+3 | | |
|---|---|---|---|---|---|---|---|---|---|---|---|---|---|---|---|---|---|---|---|---|---|---|
| | Substrates | Mean | s.e.m. | P-value | Mean | s.e.m. | P-value | Mean | s.e.m. | P-value | Mean | s.e.m. | P-value | Mean | s.e.m. | P-value | Mean | s.e.m. | P-value | Mean | s.e.m. | P-value |
| **Ac + $CO_2$** | Ac + $CO_2$ | n.d. | n.a. | n.a. | 1.06 | 0.20 | n.a. | n.d. | n.a. | n.a. | n.d. | n.a. | n.a. | 37.29 | 3.40 | n.a. | 5.00 | 0.87 | n.a. | 25.60 | 2.54 | n.a. |
| | 3Hy + $CO_2$ | 77.18 | 5.08 | n.a. | 102.13 | 1.80 | 0.00 | 22.02 | 2.55 | n.a. | 18.15 | 1.41 | n.a. | 192.84 | 4.82 | 0.00 | 104.80 | 26.56 | 0.02 | 131.94 | 29.02 | 0.02 |
| | $H_2$ + $CO_2$ | n.d. | n.a. | n.a. | n.d. | n.a. | n.a. | n.d. | n.a. | n.a. | n.d. | n.a. | n.a. | 1.30 | 0.26 | 0.00 | 1.47 | 0.08 | 0.02 | 1.55 | 0.63 | 0.00 |
| | Fe(II) + $CO_2$ | 0.67 | 0.08 | n.a. | 0.67 | 0.11 | 0.16 | 1.59 | 0.20 | n.a. | n.d. | n.a. | n.a. | 3.01 | 0.65 | 0.00 | 3.48 | 0.40 | 0.19 | n.d. | n.a. | n.a. |
| **3Hy + $CO_2$** | 3Hy + $CO_2$ | 77.18 | 5.08 | n.a. | 102.13 | 1.80 | n.a. | 22.02 | 2.55 | n.a. | 18.15 | 1.41 | n.a. | 192.84 | 4.82 | n.a. | 104.80 | 26.56 | n.a. | 131.94 | 29.02 | n.a. |
| | $H_2$ + $CO_2$ | n.a. | n.a. | n.a. | n.a. | n.a. | n.a. | n.a. | n.a. | n.a. | n.a. | n.a. | n.a. | 1.30 | 0.26 | 0.00 | 1.47 | 0.08 | 0.02 | 1.55 | 0.63 | 0.01 |
| | Fe(II) + $CO_2$ | 0.67 | 0.08 | 0.00 | 0.67 | 0.11 | 0.00 | 1.59 | 0.20 | 0.00 | n.a. | n.a. | n.a. | 3.01 | 0.65 | 0.00 | 3.48 | 0.40 | 0.02 | n.a. | n.a. | n.a. |
| **$H_2$ + $CO_2$** | $H_2$ + $CO_2$ | n.a. | n.a. | n.a. | n.a. | n.a. | n.a. | n.a. | n.a. | n.a. | n.a. | n.a. | n.a. | 1.30 | 0.26 | n.a. | 1.47 | 0.08 | n.a. | 1.55 | 0.63 | n.a. |
| | Fe(II) + $CO_2$ | 0.67 | 0.08 | n.a. | 0.67 | 0.11 | n.a. | 1.59 | 0.20 | n.a. | n.a. | n.a. | n.a. | 3.01 | 0.65 | 0.07 | 3.48 | 0.40 | 0.01 | n.a. | n.a. | n.a. |

For row 1~4, the *p*-value is between the substrate combination in the row 1 and the respective row. For row 5~7, the *p*-value is between the substrate combination in the row 5 and the respective row. For row 8~9, the *p*-value is between the substrate combination in the row 8 and the respective row.

**Supplementary Table 2-7.** *p*-value between $CO_2$ consumption/generation from different carbon and electron sources when incubated with $NH_4^+$

| | | WT+3 | | | WT+5 | | | Nif+3 | | | Nif+5 | | | Gly+3 | | | Gly+5 | | | Phb+3 | | |
|---|---|---|---|---|---|---|---|---|---|---|---|---|---|---|---|---|---|---|---|---|---|---|
| | Substrates | Mean | s.e.m. | P-value | Mean | s.e.m. | P-value | Mean | s.e.m. | P-value | Mean | s.e.m. | P-value | Mean | s.e.m. | P-value | Mean | s.e.m. | P-value | Mean | s.e.m. | P-value |
| **Ac + $CO_2$** | Ac + $CO_2$ | -59.50 | 6.67 | n.a. | -133.71 | 18.25 | n.a. | -50.53 | 8.01 | n.a. | 53.79 | 9.77 | n.a. | -197.20 | 3.50 | n.a. | -234.67 | 5.79 | n.a. | -140.01 | 42.73 | n.a. |
| | 3Hy + $CO_2$ | 75.07 | 3.34 | 0.00 | -8.94 | 2.54 | 0.01 | -13.80 | 2.67 | 0.01 | 78.67 | 15.86 | 0.25 | -29.86 | 6.52 | 0.00 | -28.50 | 9.22 | 0.00 | 15.41 | 3.07 | 0.02 |
| | $H_2$ + $CO_2$ | 176.49 | 23.17 | 0.00 | 135.17 | 13.57 | 0.00 | 88.44 | 28.81 | 0.01 | 220.27 | 10.65 | 0.00 | 29.74 | 5.77 | 0.00 | 12.15 | 2.15 | 0.00 | 141.88 | 3.42 | 0.00 |
| | Fe(II) + $CO_2$ | 147.39 | 16.91 | 0.00 | 195.99 | 17.38 | 0.00 | 273.76 | 27.25 | 0.00 | 162.04 | 10.84 | 0.00 | 74.14 | 1.29 | 0.00 | 68.73 | 4.22 | 0.00 | 113.27 | 29.70 | 0.01 |
| **3Hy + $CO_2$** | 3Hy + $CO_2$ | 75.07 | 3.34 | n.a. | -8.94 | 2.54 | n.a. | -13.80 | 2.67 | n.a. | 78.67 | 15.86 | n.a. | -29.86 | 6.52 | n.a. | -28.50 | 9.22 | n.a. | 15.41 | 3.07 | n.a. |
| | $H_2$ + $CO_2$ | 176.49 | 23.17 | 0.01 | 135.17 | 13.57 | 0.00 | 88.44 | 28.81 | 0.02 | 220.27 | 10.65 | 0.00 | 29.74 | 5.77 | 0.01 | 12.15 | 2.15 | 0.04 | 141.88 | 3.42 | 0.00 |
| | Fe(II) + $CO_2$ | 147.39 | 16.91 | 0.01 | 195.99 | 17.38 | 0.00 | 273.76 | 27.25 | 0.00 | 162.04 | 10.84 | 0.01 | 74.14 | 1.29 | 0.00 | 68.73 | 4.22 | 0.00 | 113.27 | 29.70 | 0.03 |
| **$H_2$ + $CO_2$** | $H_2$ + $CO_2$ | 176.49 | 23.17 | n.a. | 135.17 | 13.57 | n.a. | 88.44 | 28.81 | n.a. | 220.27 | 10.65 | n.a. | 29.74 | 5.77 | n.a. | 12.15 | 2.15 | n.a. | 141.88 | 3.42 | n.a. |
| | Fe(II) + $CO_2$ | 147.39 | 16.91 | 0.37 | 195.99 | 17.38 | 0.05 | 273.76 | 27.25 | 0.01 | 162.04 | 10.84 | 0.02 | 74.14 | 1.29 | 0.00 | 68.73 | 4.22 | 0.00 | 113.27 | 29.70 | 0.39 |

For row 1~4, the *p*-value is between the substrate combination in the row 1 and the respective row. For row 5~7, the *p*-value is between the substrate combination in the row 5 and the respective row. For row 8~9, the *p*-value is between the substrate combination in the row 8 and the respective row.

**Supplementary Table 2-8.** *p*-value between $CO_2$ consumption/generation from different carbon and electron sources when incubated with $N_2$

| | | WT+3 | | | WT+5 | | | Nif+3 | | | Nif+5 | | | Gly+3 | | | Gly+5 | | | Phb+3 | | |
|---|---|---|---|---|---|---|---|---|---|---|---|---|---|---|---|---|---|---|---|---|---|---|
| | Substrates | Mean | s.e.m. | P-value | Mean | s.e.m. | P-value | Mean | s.e.m. | P-value | Mean | s.e.m. | P-value | Mean | s.e.m. | P-value | Mean | s.e.m. | P-value | Mean | s.e.m. | P-value |
| **Ac + $CO_2$** | Ac + $CO_2$ | -117.57 | 20.18 | n.a. | -211.65 | 21.95 | n.a. | -47.98 | 7.77 | n.a. | 44.42 | 13.86 | n.a. | -147.61 | 9.37 | n.a. | -136.51 | 9.26 | n.a. | -218.30 | 18.13 | n.a. |
| | 3Hy + $CO_2$ | -56.86 | 10.69 | 0.06 | -41.53 | 4.57 | 0.00 | -44.87 | 10.09 | 0.82 | 35.08 | 6.72 | 0.58 | -41.12 | 8.79 | 0.00 | -114.23 | 4.52 | 0.10 | -46.65 | 6.29 | 0.00 |
| | $H_2$ + $CO_2$ | 78.52 | 13.16 | 0.00 | 87.80 | 3.30 | 0.00 | 36.41 | 2.17 | 0.00 | 141.53 | 15.73 | 0.01 | 99.04 | 15.32 | 0.00 | 61.85 | 7.62 | 0.00 | 178.17 | 28.99 | 0.00 |
| | Fe(II) + $CO_2$ | 108.44 | 13.65 | 0.00 | 45.49 | 4.73 | 0.00 | 184.23 | 10.35 | 0.00 | 78.76 | 23.25 | 0.27 | 32.65 | 8.44 | 0.00 | 9.71 | 3.19 | 0.00 | 56.99 | 6.28 | 0.00 |
| **3Hy + $CO_2$** | 3Hy + $CO_2$ | -56.86 | 10.69 | n.a. | -41.53 | 4.57 | n.a. | -44.87 | 10.09 | n.a. | 35.08 | 6.72 | n.a. | -41.12 | 8.79 | n.a. | -114.23 | 4.52 | n.a. | -46.65 | 6.29 | n.a. |
| | $H_2$ + $CO_2$ | 78.52 | 13.16 | 0.00 | 87.80 | 3.30 | 0.00 | 36.41 | 2.17 | 0.00 | 141.53 | 15.73 | 0.00 | 99.04 | 15.32 | 0.00 | 61.85 | 7.62 | 0.00 | 178.17 | 28.99 | 0.00 |
| | Fe(II) + $CO_2$ | 108.44 | 13.65 | 0.00 | 45.49 | 4.73 | 0.00 | 184.23 | 10.35 | 0.00 | 78.76 | 23.25 | 0.15 | 32.65 | 8.44 | 0.00 | 9.71 | 3.19 | 0.00 | 56.99 | 6.28 | 0.00 |
| **$H_2$ + $CO_2$** | $H_2$ + $CO_2$ | 78.52 | 13.16 | n.a. | 87.80 | 3.30 | n.a. | 36.41 | 2.17 | n.a. | 141.53 | 15.73 | n.a. | 99.04 | 15.32 | n.a. | 61.85 | 7.62 | n.a. | 178.17 | 28.99 | n.a. |
| | Fe(II) + $CO_2$ | 108.44 | 13.65 | 0.19 | 45.49 | 4.73 | 0.00 | 184.23 | 10.35 | 0.00 | 78.76 | 23.25 | 0.09 | 32.65 | 8.44 | 0.02 | 9.71 | 3.19 | 0.00 | 56.99 | 6.28 | 0.02 |

For row 1~4, the *p*-value is between the substrate combination in the row 1 and the respective row. For row 5~7, the *p*-value is between the substrate combination in the row 5 and the respective row. For row 8~9, the *p*-value is between the substrate combination in the row 8 and the respective row.

**Supplementary Table 2-9.** *p*-value between $CO_2$ consumption/generation from WT and mutants

| | | Ac-$NH_4^+$ | | | Ac-$N_2$ | | | 3Hy-$NH_4^+$ | | | 3Hy-$N_2$ | | | $H_2$-$NH_4^+$ | | | $H_2$-$N_2$ | | | Fe(II)-$NH_4^+$ | | | Fe(II)-$N_2$ | | |
|---|---|---|---|---|---|---|---|---|---|---|---|---|---|---|---|---|---|---|---|---|---|---|---|---|---|
| | Genotype | Mean | s.e.m. | P-value | Mean | s.e.m. | P-value | Mean | s.e.m. | P-value | Mean | s.e.m. | P-value | Mean | s.e.m. | P-value | Mean | s.e.m. | P-value | Mean | s.e.m. | P-value | Mean | s.e.m. | P-value |
| **3-gene** | WT | -59.50 | 6.67 | n.a. | -117.57 | 20.18 | n.a. | 75.07 | 3.34 | n.a. | -56.86 | 10.69 | n.a. | 176.49 | 23.17 | n.a. | 78.52 | 13.16 | n.a. | 147.39 | 16.91 | n.a. | 108.44 | 13.65 | n.a. |
| | Nif | -50.53 | 8.01 | 0.44 | -47.98 | 7.77 | 0.03 | -13.80 | 2.67 | 0.00 | -44.87 | 10.09 | 0.46 | 88.44 | 28.81 | 0.08 | 36.41 | 2.17 | 0.03 | 273.76 | 27.25 | 0.02 | 184.23 | 10.35 | 0.01 |
| | Gly | -197.20 | 3.50 | 0.00 | -147.61 | 9.37 | 0.25 | -29.86 | 6.52 | 0.00 | -41.12 | 8.79 | 0.32 | 29.74 | 5.77 | 0.02 | 99.04 | 15.32 | 0.37 | 74.14 | 1.29 | 0.01 | 32.65 | 8.44 | 0.01 |
| | Phb | -140.01 | 42.73 | 0.14 | -218.30 | 18.13 | 0.04 | 15.41 | 3.07 | 0.00 | -46.65 | 6.29 | 0.46 | 141.88 | 3.42 | 0.21 | 178.17 | 28.99 | 0.04 | 113.27 | 29.70 | 0.37 | 56.99 | 6.28 | 0.03 |
| **5-gene** | WT | -133.71 | 18.25 | n.a. | -211.65 | 21.95 | n.a. | -8.94 | 2.54 | n.a. | -41.53 | 4.57 | n.a. | 135.17 | 13.57 | n.a. | 87.80 | 3.30 | n.a. | 195.99 | 17.38 | n.a. | 45.49 | 4.73 | n.a. |
| | Nif | 53.79 | 9.77 | 0.00 | 44.42 | 13.86 | 0.00 | 78.67 | 15.86 | 0.02 | 35.08 | 6.72 | 0.00 | 220.27 | 10.65 | 0.01 | 141.53 | 15.73 | 0.03 | 162.04 | 10.84 | 0.17 | 78.76 | 23.25 | 0.23 |
| | Gly | -234.67 | 5.79 | 0.01 | -136.51 | 9.26 | 0.03 | -28.50 | 9.22 | 0.20 | -114.23 | 4.52 | 0.00 | 12.15 | 2.15 | 0.01 | 61.85 | 7.62 | 0.04 | 68.73 | 4.22 | 0.00 | 9.71 | 3.19 | 0.00 |

For row 1~4, the *p*-value is between the substrate combination in the row 1 and the respective row. For row 5~7, the *p*-value is between the substrate combination in the row 5 and the respective row.

**Supplementary Table 2-10.** *p*-value between carbon conversion efficiency (CCE) from WT and mutants

| | | Ac-$NH_4^+$ | | | Ac-$N_2$ | | | 3Hy-$NH_4^+$ | | | 3Hy-$N_2$ | | | $H_2$-$NH_4^+$ | | | $H_2$-$N_2$ | | | Fe(II)-$NH_4^+$ | | | Fe(II)-$N_2$ | | |
|---|---|---|---|---|---|---|---|---|---|---|---|---|---|---|---|---|---|---|---|---|---|---|---|---|---|
| | Genotype | Mean | s.e.m. | P-value | Mean | s.e.m. | P-value | Mean | s.e.m. | P-value | Mean | s.e.m. | P-value | Mean | s.e.m. | P-value | Mean | s.e.m. | P-value | Mean | s.e.m. | P-value | Mean | s.e.m. | P-value |
| **3-gene** | WT | 0.14 | 0.00 | n.a. | 0.12 | 0.02 | n.a. | 0.03 | 0.01 | n.a. | 0.13 | 0.02 | n.a. | 0.44 | 0.08 | n.a. | 1.70 | 0.32 | n.a. | 0.11 | 0.01 | n.a. | 0.15 | 0.01 | n.a. |
| | Nif | 0.25 | 0.02 | 0.01 | 1.95 | 0.26 | 0.00 | 0.37 | 0.08 | 0.01 | 0.46 | 0.07 | 0.01 | 1.61 | 0.27 | 0.01 | 4.58 | 0.23 | 0.00 | 0.14 | 0.01 | 0.03 | 0.23 | 0.01 | 0.01 |
| | Gly | 0.01 | 0.00 | 0.00 | 0.02 | 0.00 | 0.01 | 0.05 | 0.00 | 0.16 | 0.04 | 0.01 | 0.02 | 0.59 | 0.10 | 0.32 | 0.17 | 0.04 | 0.01 | n.d. | n.a. | n.a. | n.d. | n.a. | n.a. |
| | Phb | 0.06 | 0.00 | 0.00 | 0.04 | 0.00 | 0.03 | 0.03 | 0.00 | 0.40 | 0.02 | 0.00 | 0.01 | 0.07 | 0.03 | 0.01 | 0.06 | 0.01 | 0.01 | n.d. | n.a. | n.a. | 0.16 | 0.04 | 0.77 |
| **5-gene** | WT | 0.08 | 0.02 | n.a. | 0.11 | 0.03 | n.a. | 0.10 | 0.01 | n.a. | 0.24 | 0.01 | n.a. | 0.76 | 0.11 | n.a. | 1.25 | 0.01 | n.a. | 0.08 | 0.01 | n.a. | 0.27 | 0.06 | n.a. |
| | Nif | 0.47 | 0.09 | 0.01 | 1.52 | 0.30 | 0.01 | 0.49 | 0.03 | 0.00 | 0.88 | 0.16 | 0.02 | 0.82 | 0.02 | 0.61 | 0.89 | 0.21 | 0.10 | 0.18 | 0.02 | 0.01 | 0.46 | 0.12 | 0.25 |
| | Gly | n.d. | n.a. | n.a. | n.d. | n.a. | n.a. | n.d. | n.a. | n.a. | 0.03 | 0.00 | 0.00 | n.d. | n.a. | n.a. | 0.21 | 0.03 | 0.00 | n.d. | n.a. | n.a. | n.d. | n.a. | n.a. |

For row 1~4, the *p*-value is between the substrate combination in the row 1 and the respective row. For row 5~7, the *p*-value is between the substrate combination in the row 5 and the respective row.

**Supplementary Table 2-11.** *p*-value between CCE from different carbon and electron sources when incubated with $NH_4^+$

| | | WT+3 | | | WT+5 | | | Nif+3 | | | Nif+5 | | | Gly+3 | | | Gly+5 | | | Phb+3 | | |
|---|---|---|---|---|---|---|---|---|---|---|---|---|---|---|---|---|---|---|---|---|---|---|
| | Substrates | Mean | s.e.m. | P-value | Mean | s.e.m. | P-value | Mean | s.e.m. | P-value | Mean | s.e.m. | P-value | Mean | s.e.m. | P-value | Mean | s.e.m. | P-value | Mean | s.e.m. | P-value |
| **Ac + $CO_2$** | Ac + $CO_2$ | 0.14 | 0.00 | n.a. | 0.08 | 0.02 | n.a. | 0.25 | 0.02 | n.a. | 0.47 | 0.09 | n.a. | 0.01 | 0.00 | n.a. | n.d. | n.a. | n.a. | 0.06 | 0.00 | n.a. |
| | 3Hy + $CO_2$ | 0.03 | 0.01 | 0.00 | 0.10 | 0.01 | 0.41 | 0.37 | 0.08 | 0.20 | 0.49 | 0.03 | 0.83 | 0.05 | 0.00 | 0.00 | n.d. | n.a. | n.a. | 0.03 | 0.00 | 0.00 |
| | $H_2$ + $CO_2$ | 0.44 | 0.08 | 0.02 | 0.76 | 0.11 | 0.00 | 1.61 | 0.27 | 0.01 | 0.82 | 0.02 | 0.01 | 0.59 | 0.10 | 0.00 | n.d. | n.a. | n.a. | 0.07 | 0.03 | 0.61 |
| | Fe(II) + $CO_2$ | 0.11 | 0.01 | 0.03 | 0.08 | 0.01 | 0.93 | 0.14 | 0.01 | 0.01 | 0.18 | 0.02 | 0.03 | n.d. | n.a. | n.a. | n.d. | n.a. | n.a. | n.d. | n.a. | n.a. |
| **3Hy + $CO_2$** | 3Hy + $CO_2$ | 0.03 | 0.01 | n.a. | 0.10 | 0.01 | n.a. | 0.37 | 0.08 | n.a. | 0.49 | 0.03 | n.a. | 0.05 | 0.00 | n.a. | n.a. | n.a. | n.a. | 0.03 | 0.00 | n.a. |
| | $H_2$ + $CO_2$ | 0.44 | 0.08 | 0.01 | 0.76 | 0.11 | 0.00 | 1.61 | 0.27 | 0.01 | 0.82 | 0.02 | 0.00 | 0.59 | 0.10 | 0.03 | n.a. | n.a. | n.a. | 0.07 | 0.03 | 0.15 |
| | Fe(II) + $CO_2$ | 0.11 | 0.01 | 0.00 | 0.08 | 0.01 | 0.25 | 0.14 | 0.01 | 0.04 | 0.18 | 0.02 | 0.00 | n.a. | n.a. | n.a. | n.a. | n.a. | n.a. | n.a. | n.a. | n.a. |
| **$H_2$ + $CO_2$** | $H_2$ + $CO_2$ | 0.44 | 0.08 | n.a. | 0.76 | 0.11 | n.a. | 1.61 | 0.27 | n.a. | 0.82 | 0.02 | n.a. | 0.59 | 0.10 | n.a. | n.a. | n.a. | n.a. | 0.07 | 0.03 | n.a. |
| | Fe(II) + $CO_2$ | 0.11 | 0.01 | 0.01 | 0.08 | 0.01 | 0.00 | 0.14 | 0.01 | 0.01 | 0.18 | 0.02 | 0.00 | n.a. | n.a. | n.a. | n.a. | n.a. | n.a. | n.a. | n.a. | n.a. |

For row 1~4, the *p*-value is between the substrate combination in the row 1 and the respective row. For row 5~7, the *p*-value is between the substrate combination in the row 5 and the respective row. For row 8~9, the *p*-value is between the substrate combination in the row 8 and the respective row.

**Supplementary Table 2-12.** *p*-value between CCE from different carbon and electron sources when incubated with $N_2$

| | | WT+3 | | | WT+5 | | | Nif+3 | | | Nif+5 | | | Gly+3 | | | Gly+5 | | | Phb+3 | | |
|---|---|---|---|---|---|---|---|---|---|---|---|---|---|---|---|---|---|---|---|---|---|---|
| | Substrates | Mean | s.e.m. | P-value | Mean | s.e.m. | P-value | Mean | s.e.m. | P-value | Mean | s.e.m. | P-value | Mean | s.e.m. | P-value | Mean | s.e.m. | P-value | Mean | s.e.m. | P-value |
| **Ac + $CO_2$** | Ac + $CO_2$ | 0.12 | 0.02 | n.a. | 0.11 | 0.03 | n.a. | 1.95 | 0.26 | n.a. | 1.52 | 0.30 | n.a. | 0.02 | 0.00 | n.a. | n.d. | n.a. | n.a. | 0.04 | 0.00 | n.a. |
| | 3Hy + $CO_2$ | 0.13 | 0.02 | 0.87 | 0.24 | 0.01 | 0.02 | 0.46 | 0.07 | 0.00 | 0.88 | 0.16 | 0.13 | 0.04 | 0.01 | 0.05 | 0.03 | 0.00 | n.a. | 0.02 | 0.00 | 0.02 |
| | $H_2$ + $CO_2$ | 1.70 | 0.32 | 0.01 | 1.25 | 0.01 | 0.00 | 4.58 | 0.23 | 0.00 | 0.89 | 0.21 | 0.23 | 0.17 | 0.04 | 0.02 | 0.21 | 0.03 | n.a. | 0.06 | 0.01 | 0.11 |
| | Fe(II) + $CO_2$ | 0.15 | 0.01 | 0.37 | 0.27 | 0.06 | 0.08 | 0.23 | 0.01 | 0.00 | 0.46 | 0.12 | 0.03 | n.d. | n.a. | n.a. | n.d. | n.a. | n.a. | 0.16 | 0.04 | 0.03 |
| **3Hy + $CO_2$** | 3Hy + $CO_2$ | 0.13 | 0.02 | n.a. | 0.24 | 0.01 | n.a. | 0.46 | 0.07 | n.a. | 0.88 | 0.16 | n.a. | 0.04 | 0.01 | n.a. | 0.03 | 0.00 | n.a. | 0.02 | 0.00 | n.a. |
| | $H_2$ + $CO_2$ | 1.70 | 0.32 | 0.01 | 1.25 | 0.01 | 0.00 | 4.58 | 0.23 | 0.00 | 0.89 | 0.21 | 0.98 | 0.17 | 0.04 | 0.03 | 0.21 | 0.03 | 0.00 | 0.06 | 0.01 | 0.02 |
| | Fe(II) + $CO_2$ | 0.15 | 0.01 | 0.41 | 0.27 | 0.06 | 0.59 | 0.23 | 0.01 | 0.03 | 0.46 | 0.12 | 0.11 | n.a. | n.a. | n.a. | n.a. | n.a. | n.a. | 0.16 | 0.04 | 0.02 |
| **$H_2$ + $CO_2$** | $H_2$ + $CO_2$ | 1.70 | 0.32 | n.a. | 1.25 | 0.01 | n.a. | 4.58 | 0.23 | n.a. | 0.89 | 0.21 | n.a. | 0.17 | 0.04 | n.a. | 0.21 | 0.03 | n.a. | 0.06 | 0.01 | n.a. |
| | Fe(II) + $CO_2$ | 0.15 | 0.01 | 0.01 | 0.27 | 0.06 | 0.00 | 0.23 | 0.01 | 0.00 | 0.46 | 0.12 | 0.15 | n.a. | n.a. | n.a. | n.a. | n.a. | n.a. | 0.16 | 0.04 | 0.06 |

For row 1~4, the *p*-value is between the substrate combination in the row 1 and the respective row. For row 5~7, the *p*-value is between the substrate combination in the row 5 and the respective row. For row 8~9, the *p*-value is between the substrate combination in the row 8 and the respective row.

**Supplementary Table 2-13.** *p*-value between CCE production from different nitrogen sources

| | | WT+3 | | | WT+5 | | | Nif+3 | | | Nif+5 | | | Gly+3 | | | Gly+5 | | | Phb+3 | | |
|---|---|---|---|---|---|---|---|---|---|---|---|---|---|---|---|---|---|---|---|---|---|---|
| | N source | Mean | s.e.m. | P-value | Mean | s.e.m. | P-value | Mean | s.e.m. | P-value | Mean | s.e.m. | P-value | Mean | s.e.m. | P-value | Mean | s.e.m. | P-value | Mean | s.e.m. | P-value |
| **Ac + $CO_2$** | $NH_4^+$ | 0.14 | 0.00 | 0.66 | 0.08 | 0.02 | 0.52 | 0.25 | 0.02 | 0.00 | 0.47 | 0.09 | 0.03 | 0.01 | 0.00 | 0.03 | n.d. | n.a. | n.a. | 0.06 | 0.00 | 0.05 |
| | $N_2$ | 0.12 | 0.02 | | 0.11 | 0.03 | | 1.95 | 0.26 | | 1.52 | 0.30 | | 0.02 | 0.00 | | n.d. | n.a. | | 0.04 | 0.00 | |
| **3Hy + $CO_2$** | $NH_4^+$ | 0.03 | 0.01 | 0.01 | 0.10 | 0.01 | 0.00 | 0.37 | 0.08 | 0.45 | 0.49 | 0.03 | 0.08 | 0.05 | 0.00 | 0.63 | n.d. | n.a. | n.a. | 0.03 | 0.00 | 0.10 |
| | $N_2$ | 0.13 | 0.02 | | 0.24 | 0.01 | | 0.46 | 0.07 | | 0.88 | 0.16 | | 0.04 | 0.01 | | 0.03 | 0.00 | | 0.02 | 0.00 | |
| **$H_2$ + $CO_2$** | $NH_4^+$ | 0.44 | 0.08 | 0.02 | 0.76 | 0.11 | 0.01 | 1.61 | 0.27 | 0.00 | 0.82 | 0.02 | 0.72 | 0.59 | 0.10 | 0.02 | n.d. | n.a. | n.a. | 0.07 | 0.03 | 0.64 |
| | $N_2$ | 1.70 | 0.32 | | 1.25 | 0.01 | | 4.58 | 0.23 | | 0.89 | 0.21 | | 0.17 | 0.04 | | 0.21 | 0.03 | | 0.06 | 0.01 | |
| **Fe(II) + $CO_2$** | $NH_4^+$ | 0.11 | 0.01 | 0.04 | 0.08 | 0.01 | 0.04 | 0.14 | 0.01 | 0.00 | 0.18 | 0.02 | 0.09 | n.d. | n.a. | n.a. | n.d. | n.a. | n.a. | n.d. | n.a. | n.a. |
| | $N_2$ | 0.15 | 0.01 | | 0.27 | 0.06 | | 0.23 | 0.01 | | 0.46 | 0.12 | | n.d. | n.a. | | n.d. | n.a. | | 0.16 | 0.04 | |

**Supplementary Table 2-14.** *p*-value between electron conversion efficiency from different carbon and electron sources when incubated with $NH_4^+$

| | | WT+3 | | | WT+5 | | | Nif+3 | | | Nif+5 | | | Gly+3 | | | Gly+5 | | | Phb+3 | | |
|---|---|---|---|---|---|---|---|---|---|---|---|---|---|---|---|---|---|---|---|---|---|---|
| | Substrates | Mean | s.e.m. | P-value | Mean | s.e.m. | P-value | Mean | s.e.m. | P-value | Mean | s.e.m. | P-value | Mean | s.e.m. | P-value | Mean | s.e.m. | P-value | Mean | s.e.m. | P-value |
| **Ac + $CO_2$** | Ac + $CO_2$ | 0.03 | 0.00 | n.a. | 0.02 | 0.00 | n.a. | 0.06 | 0.01 | n.a. | 0.12 | 0.03 | n.a. | 0.00 | 0.00 | n.a. | n.d. | n.a. | n.a. | 0.01 | 0.00 | n.a. |
| | 3Hy + $CO_2$ | 0.00 | 0.00 | 0.00 | 0.01 | 0.00 | 0.06 | 0.03 | 0.01 | 0.02 | 0.04 | 0.00 | 0.07 | 0.00 | 0.00 | 0.02 | n.d. | n.a. | n.a. | 0.00 | 0.00 | 0.00 |
| | $H_2$ + $CO_2$ | 0.43 | 0.09 | 0.01 | 0.59 | 0.14 | 0.02 | 0.57 | 0.08 | 0.00 | 0.55 | 0.01 | 0.00 | 0.05 | 0.01 | 0.00 | n.d. | n.a. | n.a. | 0.03 | 0.01 | 0.30 |
| | Fe(II) + $CO_2$ | 2.25 | 0.35 | 0.00 | 6.45 | 1.73 | 0.02 | 5.82 | 0.30 | 0.00 | 4.41 | 0.49 | 0.00 | n.d. | n.a. | n.a. | n.d. | n.a. | n.a. | n.d. | n.a. | n.a. |
| **3Hy + $CO_2$** | 3Hy + $CO_2$ | 0.00 | 0.00 | n.a. | 0.01 | 0.00 | n.a. | 0.03 | 0.01 | n.a. | 0.04 | 0.00 | n.a. | 0.00 | 0.00 | n.a. | n.a. | n.a. | n.a. | 0.00 | 0.00 | n.a. |
| | $H_2$ + $CO_2$ | 0.43 | 0.09 | 0.01 | 0.59 | 0.14 | 0.01 | 0.57 | 0.08 | 0.00 | 0.55 | 0.01 | 0.00 | 0.05 | 0.01 | 0.01 | n.a. | n.a. | n.a. | 0.03 | 0.01 | 0.06 |
| | Fe(II) + $CO_2$ | 2.25 | 0.35 | 0.00 | 6.45 | 1.73 | 0.02 | 5.82 | 0.30 | 0.00 | 4.41 | 0.49 | 0.00 | n.a. | n.a. | n.a. | n.a. | n.a. | n.a. | n.a. | n.a. | n.a. |
| **$H_2$ + $CO_2$** | $H_2$ + $CO_2$ | 0.43 | 0.09 | n.a. | 0.59 | 0.14 | n.a. | 0.57 | 0.08 | n.a. | 0.55 | 0.01 | n.a. | 0.05 | 0.01 | n.a. | n.a. | n.a. | n.a. | 0.03 | 0.01 | n.a. |
| | Fe(II) + $CO_2$ | 2.25 | 0.35 | 0.01 | 6.45 | 1.73 | 0.03 | 5.82 | 0.30 | 0.00 | 4.41 | 0.49 | 0.00 | n.a. | n.a. | n.a. | n.a. | n.a. | n.a. | n.a. | n.a. | n.a. |

For row 1~4, the *p*-value is between the substrate combination in the row 1 and the respective row. For row 5~7, the *p*-value is between the substrate combination in the row 5 and the respective row. For row 8~9, the *p*-value is between the substrate combination in the row 8 and the respective row.

**Supplementary Table 2-15.** *p*-value between electron conversion efficiency from different carbon and electron sources when incubated with $N_2$

| | | WT+3 | | | WT+5 | | | Nif+3 | | | Nif+5 | | | Gly+3 | | | Gly+5 | | | Phb+3 | | |
|---|---|---|---|---|---|---|---|---|---|---|---|---|---|---|---|---|---|---|---|---|---|---|
| | Substrates | Mean | s.e.m. | P-value | Mean | s.e.m. | P-value | Mean | s.e.m. | P-value | Mean | s.e.m. | P-value | Mean | s.e.m. | P-value | Mean | s.e.m. | P-value | Mean | s.e.m. | P-value |
| **Ac + $CO_2$** | Ac + $CO_2$ | 0.03 | 0.01 | n.a. | 0.03 | 0.01 | n.a. | 0.49 | 0.06 | n.a. | 0.38 | 0.07 | n.a. | 0.00 | 0.00 | n.a. | n.d. | n.a. | n.a. | 0.01 | 0.00 | n.a. |
| | 3Hy + $CO_2$ | 0.01 | 0.00 | 0.03 | 0.02 | 0.00 | 0.42 | 0.04 | 0.01 | 0.00 | 0.07 | 0.01 | 0.02 | 0.00 | 0.00 | 0.21 | 0.00 | 0.00 | n.a. | 0.00 | 0.00 | 0.00 |
| | $H_2$ + $CO_2$ | 0.42 | 0.03 | 0.00 | 0.32 | 0.01 | 0.00 | 0.50 | 0.04 | 0.91 | 0.49 | 0.04 | 0.36 | 0.04 | 0.00 | 0.00 | 0.03 | 0.00 | n.a. | 0.03 | 0.00 | 0.02 |
| | Fe(II) + $CO_2$ | 2.27 | 0.39 | 0.00 | 2.52 | 0.31 | 0.00 | 10.37 | 0.91 | 0.00 | 12.47 | 1.37 | 0.00 | n.d. | n.a. | n.a. | n.d. | n.a. | n.a. | 0.02 | 0.01 | 0.14 |
| **3Hy + $CO_2$** | 3Hy + $CO_2$ | 0.01 | 0.00 | n.a. | 0.02 | 0.00 | n.a. | 0.04 | 0.01 | n.a. | 0.07 | 0.01 | n.a. | 0.00 | 0.00 | n.a. | 0.00 | 0.00 | n.a. | 0.00 | 0.00 | n.a. |
| | $H_2$ + $CO_2$ | 0.42 | 0.03 | 0.00 | 0.32 | 0.01 | 0.00 | 0.50 | 0.04 | 0.00 | 0.49 | 0.04 | 0.00 | 0.04 | 0.00 | 0.00 | 0.03 | 0.00 | 0.00 | 0.03 | 0.00 | 0.01 |
| | Fe(II) + $CO_2$ | 2.27 | 0.39 | 0.00 | 2.52 | 0.31 | 0.00 | 10.37 | 0.91 | 0.00 | 12.47 | 1.37 | 0.00 | n.a. | n.a. | n.a. | n.a. | n.a. | n.a. | 0.02 | 0.01 | 0.03 |
| **$H_2$ + $CO_2$** | $H_2$ + $CO_2$ | 0.42 | 0.03 | n.a. | 0.32 | 0.01 | n.a. | 0.50 | 0.04 | n.a. | 0.49 | 0.04 | n.a. | 0.04 | 0.00 | n.a. | 0.03 | 0.00 | n.a. | 0.03 | 0.00 | n.a. |
| | Fe(II) + $CO_2$ | 2.27 | 0.39 | 0.01 | 2.52 | 0.31 | 0.00 | 10.37 | 0.91 | 0.00 | 12.47 | 1.37 | 0.01 | n.a. | n.a. | n.a. | n.a. | n.a. | n.a. | 0.02 | 0.01 | 0.43 |

For row 1~4, the *p*-value is between the substrate combination in the row 1 and the respective row. For row 5~7, the *p*-value is between the substrate combination in the row 5 and the respective row. For row 8~9, the *p*-value is between the substrate combination in the row 8 and the respective row.

**Supplementary Table 2-16.** *p*-value between electron conversion efficiency from WT and mutants

| | | Ac-$NH_4^+$ | | | Ac-$N_2$ | | | 3Hy-$NH_4^+$ | | | 3Hy-$N_2$ | | | $H_2$-$NH_4^+$ | | | $H_2$-$N_2$ | | | Fe(II)-$NH_4^+$ | | | Fe(II)-$N_2$ | | |
|---|---|---|---|---|---|---|---|---|---|---|---|---|---|---|---|---|---|---|---|---|---|---|---|---|---|
| | Genotype | Mean | s.e.m. | P-value | Mean | s.e.m. | P-value | Mean | s.e.m. | P-value | Mean | s.e.m. | P-value | Mean | s.e.m. | P-value | Mean | s.e.m. | P-value | Mean | s.e.m. | P-value | Mean | s.e.m. | P-value |
| **3-gene** | WT | 0.03 | 0.00 | n.a. | 0.03 | 0.01 | n.a. | 0.00 | 0.00 | n.a. | 0.01 | 0.00 | n.a. | 0.43 | 0.09 | n.a. | 0.42 | 0.03 | n.a. | 2.25 | 0.35 | n.a. | 2.27 | 0.39 | n.a. |
| | Nif | 0.06 | 0.01 | 0.01 | 0.49 | 0.06 | 0.00 | 0.03 | 0.01 | 0.01 | 0.04 | 0.01 | 0.01 | 0.57 | 0.08 | 0.32 | 0.50 | 0.04 | 0.22 | 5.82 | 0.30 | 0.00 | 10.37 | 0.91 | 0.00 |
| | Gly | 0.00 | 0.00 | 0.00 | 0.00 | 0.00 | 0.01 | 0.00 | 0.00 | 0.21 | 0.00 | 0.00 | 0.02 | 0.05 | 0.01 | 0.01 | 0.04 | 0.00 | 0.00 | n.d. | n.a. | n.a. | n.d. | n.a. | n.a. |
| | Phb | 0.01 | 0.00 | 0.00 | 0.01 | 0.00 | 0.03 | 0.00 | 0.00 | 0.32 | 0.00 | 0.00 | 0.01 | 0.03 | 0.01 | 0.01 | 0.03 | 0.00 | 0.00 | n.d. | n.a. | n.a. | 0.02 | 0.01 | 0.00 |
| **5-gene** | WT | 0.02 | 0.00 | n.a. | 0.03 | 0.01 | n.a. | 0.01 | 0.00 | n.a. | 0.02 | 0.00 | n.a. | 0.59 | 0.14 | n.a. | 0.32 | 0.01 | n.a. | 6.45 | 1.73 | n.a. | 2.52 | 0.31 | n.a. |
| | Nif | 0.12 | 0.03 | 0.04 | 0.38 | 0.07 | 0.01 | 0.04 | 0.00 | 0.00 | 0.07 | 0.01 | 0.02 | 0.55 | 0.01 | 0.79 | 0.49 | 0.04 | 0.02 | 4.41 | 0.49 | 0.32 | 12.47 | 1.37 | 0.00 |
| | Gly | n.d. | n.a. | n.a. | n.d. | n.a. | n.a. | n.d. | n.a. | n.a. | 0.00 | 0.00 | 0.00 | n.d. | n.a. | n.a. | 0.03 | 0.00 | 0.00 | n.d. | n.a. | n.a. | n.d. | n.a. | n.a. |

For row 1~4, the *p*-value is between the substrate combination in the row 1 and the respective row. For row 5~7, the *p*-value is between the substrate combination in the row 5 and the respective row.

**Supplementary Table 2-17.** *p*-value between from different electricity sources during photoelectroautotrophy

a. *n*-butanol, acetone, CCE, electron conversion efficiency, electrical energy conversion efficiency (EECE)

| | ***n*-Butanol (mg/L)** | | | **Acetone mg/L** | | | **CCE (%)** | | | **Electron conversion efficiency (%)** | | | **EECE (%)** | | |
|---|---|---|---|---|---|---|---|---|---|---|---|---|---|---|---|
| Platform | Mean | s.e.m. | P-value | Mean | s.e.m. | P-value | Mean | s.e.m. | P-value | Mean | s.e.m. | P-value | Mean | s.e.m. | P-value |
| SO-HA-$NH_4^+$ | 0.50 | 0.05 | 0.20 | 0.50 | 0.07 | 0.00 | 0.06 | 0.01 | 0.01 | 1.79 | 0.14 | 0.00 | 4.80 | 0.38 | 0.00 |
| PO-HA-$NH_4^+$ | 0.32 | 0.08 | | 0.00 | 0.00 | | 0.18 | 0.01 | | 43.93 | 2.01 | | 117.53 | 5.37 | |
| SO-IR-$NH_4^+$ | 0.58 | 0.07 | 0.01 | 1.93 | 0.08 | 0.00 | 0.07 | 0.01 | 0.18 | 1.68 | 0.13 | 0.00 | 4.50 | 0.34 | 0.00 |
| PO-IR-$NH_4^+$ | 0.63 | 0.02 | | 1.21 | 0.13 | | 0.49 | 0.06 | | 9.91 | 1.07 | | 26.52 | 2.87 | |
| SO-HA-$N_2$ | 0.91 | 0.07 | 0.56 | 1.25 | 0.08 | 0.04 | 0.11 | 0.01 | 0.00 | 3.57 | 0.13 | 0.02 | 9.55 | 0.34 | 0.02 |
| PO-HA-$N_2$ | 0.33 | 0.07 | | 0.00 | 0.00 | | 0.10 | 0.00 | | 49.02 | 1.48 | | 131.14 | 3.97 | |
| SO-IR-$N_2$ | 0.19 | 0.02 | 0.11 | 0.55 | 0.09 | 0.05 | 0.02 | 0.00 | 0.03 | 0.60 | 0.07 | 0.01 | 1.62 | 0.20 | 0.00 |
| PO-IR-$N_2$ | 0.46 | 0.09 | | 1.67 | 0.24 | | 0.40 | 0.06 | | 6.28 | 0.64 | | 16.62 | 1.01 | |

b. $CO_2$ consumption, electron uptake, percentage of alive cell on electrode, total number of cells on electrode

| | $CO_2$ consumption (μmol) | | | Electron up take (C) | | | Alive cell percentatge (%) | | | Total number of cell | | |
|---|---|---|---|---|---|---|---|---|---|---|---|---|
| Platform | Mean | s.e.m. | P-value | Mean | s.e.m. | P-value | Mean | s.e.m. | P-value | Mean | s.e.m. | P-value |
| SO-HA-$NH_4$ | 1775.8435 | 77.3031 | 0.0001 | 69.1375 | 3.0865 | 0.0011 | 0.4391 | 0.0500 | 0.6690 | 1596.2500 | 80.2566 | 0.0384 |
| PO-HA-$NH_4$ | 596.0338 | 91.5679 | | 1.8649 | 0.7296 | | 0.4102 | 0.0653 | | 670.0000 | 251.7300 | |
| SO-IR-$NH_4$ | 2019.5304 | 23.9546 | 0.0046 | 85.7635 | 5.7810 | 0.0002 | 0.3648 | 0.0355 | 0.4306 | 2967.4444 | 240.9069 | 0.4477 |
| PO-IR-$NH_4$ | 486.7531 | 152.5035 | | 13.1370 | 5.7737 | | 0.4447 | 0.0324 | | 768.0000 | 68.3537 | |
| SO-HA-$N_2$ | 1790.8779 | 18.5053 | 0.0001 | 63.9758 | 1.2722 | 0.0008 | 0.5210 | 0.0375 | 0.0851 | 2200.0000 | 210.2464 | 0.0012 |
| PO-HA-$N_2$ | 602.5003 | 207.6853 | | 1.6758 | 0.4467 | | 0.4947 | 0.0145 | | 1677.5000 | 760.1398 | |
| SO-IR-$N_2$ | 1947.8308 | 102.5876 | 0.0002 | 80.3773 | 0.2595 | 0.0003 | 0.4472 | 0.0368 | 0.0366 | 1310.4444 | 452.2298 | 0.0805 |
| PO-HA-$N_2$ | 602.5003 | 207.6853 | | 1.6758 | 0.4467 | | 0.4947 | 0.0145 | | 1677.5000 | 760.1398 | |

**Supplementary Table 2-18.** *p*-value between different light sources during photoelectroautotrophy

a. *n*-butanol, acetone, CCE, electron conversion efficiency, EECE

| | ***n*-Butanol (mg/L)** | | | **Acetone mg/L** | | | **CCE (%)** | | | **Electron conversion efficiency (%)** | | | **EECE (%)** | | |
|---|---|---|---|---|---|---|---|---|---|---|---|---|---|---|---|
| Platform | Mean | s.e.m. | P-value | Mean | s.e.m. | P-value | Mean | s.e.m. | P-value | Mean | s.e.m. | P-value | Mean | s.e.m. | P-value |
| SO-HA-$NH_4^+$ | 0.50 | 0.05 | 0.46 | 0.50 | 0.07 | 0.01 | 0.06 | 0.01 | 0.56 | 1.79 | 0.14 | 0.61 | 4.80 | 0.38 | 0.61 |
| SO-IR-$NH_4^+$ | 0.58 | 0.07 | | 1.93 | 0.08 | | 0.07 | 0.01 | | 1.68 | 0.13 | | 4.50 | 0.34 | |
| SO-HA-$N_2$ | 0.91 | 0.00 | 0.00 | 1.25 | 0.01 | 0.01 | 0.11 | 0.00 | 0.00 | 3.57 | 0.03 | 0.00 | 9.55 | 0.09 | 0.00 |
| SO-IR-$N_2$ | 0.19 | 0.02 | | 0.55 | 0.09 | | 0.02 | 0.00 | | 0.60 | 0.07 | | 1.62 | 0.20 | |
| PO-HA-$NH_4^+$ | 0.32 | 0.08 | 0.06 | 0.00 | 0.00 | 0.00 | 0.18 | 0.01 | 0.04 | 43.93 | 2.01 | 0.00 | 117.53 | 5.37 | 0.00 |
| PO-IR-$NH_4^+$ | 0.63 | 0.02 | | 1.21 | 0.13 | | 0.49 | 0.06 | | 9.91 | 1.07 | | 26.52 | 2.87 | |
| PO-HA-$N_2$ | 0.33 | 0.07 | 0.38 | 0.00 | 0.00 | 0.00 | 0.10 | 0.00 | 0.04 | 49.02 | 1.48 | 0.00 | 131.14 | 3.97 | 0.00 |
| PO-HA-$N_2$ | 0.46 | 0.09 | | 1.67 | 0.24 | | 0.40 | 0.06 | | 6.28 | 0.64 | | 16.62 | 1.01 | |

b. $CO_2$ consumption, electron uptake, percentage of alive cell on electrode, total number of cells on electrode

| | $CO_2$ consumption (µmol) | | | Electron up take (C) | | | Alive cell percentatge (%) | | | Total number of cell | | |
|---|---|---|---|---|---|---|---|---|---|---|---|---|
| Platform | Mean | s.e.m. | P-value | Mean | s.e.m. | P-value | Mean | s.e.m. | P-value | Mean | s.e.m. | P-value |
| SO-HA-$NH_4$ | 1775.8435 | 77.3031 | 0.0064 | 69.1375 | 3.0865 | 0.0697 | 0.4391 | 0.0500 | 0.1410 | 1596.2500 | 80.2566 | 0.0050 |
| SO-IR-$NH_4$ | 2019.5304 | 23.9546 | | 85.7635 | 5.7810 | | 0.3648 | 0.0355 | | 2967.4444 | 240.9069 | |
| SO-HA-$N_2$ | 1790.8779 | 18.5053 | 0.1345 | 63.9758 | 1.2722 | 0.0031 | 0.5210 | 0.0375 | 0.0719 | 2200.0000 | 210.2464 | 0.0872 |
| SO-IR-$N_2$ | 1947.8308 | 102.5876 | | 80.3773 | 0.2595 | | 0.4472 | 0.0368 | | 1310.4444 | 452.2298 | |
| PO-HA-$NH_4$ | 596.0338 | 91.5679 | 0.3473 | 1.8649 | 0.7296 | 0.0798 | 0.4102 | 0.0653 | 0.5727 | 670.0000 | 251.7300 | 0.6483 |
| PO-IR-$NH_4$ | 486.7531 | 152.5035 | | 13.1370 | 5.7737 | | 0.4447 | 0.0324 | | 768.0000 | 68.3537 | |
| PO-HA-$N_2$ | 602.5003 | 207.6853 | 0.1063 | 1.6758 | 0.4467 | 0.0292 | 0.4947 | 0.0145 | 0.1998 | 1677.5000 | 760.1398 | 0.4994 |
| PO-HA-$N_2$ | 248.1555 | 13.3554 | | 15.3200 | 4.6405 | | 0.5520 | 0.0458 | | 2022.5556 | 276.2200 | |

All *p*-values are calculated using mean and s.e.m. (standard error of mean) of at three (values from photoelectroautotrophy has only two biological replicates) biological replicates. Ac: acetate; 3Hy: 3-hydroxybutyrate; $H_2$: hydrogen; Fe(II): ferrous iron; $CO_2$: carbon dioxide; $NH_4$: ammonium; $N_2$: dinitrogen gas; SO: solar panel; PO: potentiostat; HA: halogen light; IR: infrared light. Nif: Δ*nifA*1Δ*nifA*2; Gly: Δ*glgA*1; Phb: Δ*phaC*1Δ*phaC*2. WT+3: WT with 3-gene cassette; WT+5: WT with 5-gene cassette; Nif+3: *ΔnifA1ΔnifA2* with 3-gene cassette; Nif+5: *ΔnifA1ΔnifA2* with 5-gene cassette; Gly+3: *ΔglgA* with 3-gene cassette; Gly+5: *ΔglgA* with 5-gene cassette; Phb+3: *ΔphaC1ΔphaC2* with 3-gene cassette; n.d.: non-detectable; n. a.: non-applicable due to non-detectable results

**Supplementary Table 3. Growth study of TIE-1 and various mutants under different conditions.**

**Supplementary Table 3-1 WT with pAB423 in media supplied with deferent amount of *n*-butanol.**

| **Concentration of *n*-Butanol (mg/L)** | 0 | 2025 | 4050 | 8100 | 16200 |
|---|---|---|---|---|---|
| **Doubling time (hr)** | 15.00 | 20.091 | no growth | no growth | no growth |

**Supplementary Table 3-2 WT with pAB423 in media supplied with deferent amount of acetone.**

| **Concentration of Acetone (mg/L)** | 0 | 1960 | 3920 | 7840 | 15680 |
|---|---|---|---|---|---|
| **Doubling time (hr)** | 7.30 | 6.74 | 5.82 | 5.56 | 4.88 |

**Supplementary Table 4. Specific oxidation state and number of electrons required per mole of *n*-butanol.**

| **Carbon source** | **The oxidation state of carbon in carbon source** | **The oxidation state of carbon in *n*-butanol** | **Mole carbon needed per mole *n*-butanol** | **Electron needed per mole *n*-butanol** |
|---|---|---|---|---|
| Acetate | 0 | -2 | 4 | 8 |
| 3-hydroxybutyrate | -0.5 | -2 | 4 | 6 |
| Carbon dioxide | 4 | -2 | 4 | 24 |
